# The macrophage genetic cassette inr/dtor/pvf2 is a nutritional status checkpoint for developmental timing

**DOI:** 10.1101/2023.01.05.522883

**Authors:** Sergio Juarez-Carreño, Frederic Geissmann

## Abstract

A small number of signaling molecules, used reiteratively, control differentiation programs, but the mechanisms that adapt developmental timing to environmental cues are less understood. We report here that a macrophage *inr/dtor/pvf2* genetic cassette is a developmental timing checkpoint in *Drosophila*, which either licenses or delays biosynthesis of the steroid hormone in the endocrine gland and metamorphosis according to the larval nutritional status. Insulin-Receptor/dTor signaling in macrophages is required and sufficient for production of the PDGF/VEGF family growth factor Pvf2, which turns on transcription of the sterol biosynthesis Halloween genes in the prothoracic gland via its receptor Pvr. In response to a starvation event or genetic manipulation, low Pvf2 signal delays steroid biosynthesis until it becomes Pvr-independent, thereby prolonging larval growth before pupation. The significance of this developmental timing checkpoint for host fitness is illustrated by the observation that it regulates the size of the pupae and adult flies.

**One sentence summary:** A macrophage Inr/dtor/pvf2 cassette is a nutritional status checkpoint that controls developmental timing via steroid hormone biosynthesis in the endocrine gland.

## Introduction

Controlled developmental timing is essential for fitness in the plant and animal phyla, because it allows the success of critical transitions such as sexual reproduction through flowering in plants [1], puberty/sexual maturation in mammals [2, 3], and metamorphosis from larvae to adults in insects [4, 5]. Puberty and metamorphosis are triggered by the production of steroid hormones in endocrine glands. In mammals the hypothalamus gonadotropin-releasing hormone (GnRH) controls steroid synthesis via the anterior pituitary hormones (ACTH, LH, FSH) [2, 3, 6]. In *Drosophila* the GnRH superfamily hormone Corazonin (Crz), controls steroid synthesis via the prothoracicotropic hormone (PTTH) produced by the PTTH neurons [4, 7-9]. Although genetic and molecular studies have identified the limited number of conserved signaling molecules that are used reiteratively to control puberty, metamorphosis, and more generally differentiation [10], the mechanisms that adapt developmental timing to external cues are less understood. Nutritional status is an important factor that can delay development, when sub-optimal nutrition could compromise the fitness of the adult. Specifically, the production of fat-derived signals (FDS) acting on the brain, such as Leptin in mammals, and its *Drosophila* ortholog Upd2, with Eiger and Stu, represents critical checkpoints for nutritional status that control puberty and metamorphosis [8, 11-17]. The developmental timing of puberty and metamorphosis therefore integrates nutritional quality control checkpoints, which rely on inter-organ communication networks, to optimize the timing of steroid hormone synthesis and the fitness of adults. However, these inter-organ and intercellular communication networks remain poorly understood. Macrophages are accessory cells present throughout tissues, and recent findings have attracted attention to their roles in developmental processes, inter-organ communication, and metabolism including in response to nutrient availability [18-24] and steroid hormone synthesis [25, 26]. Moreover, macrophage deficiency delays metamorphosis in *Drosophila* [27], and impairs steroid hormone synthesis and sexual maturation of rodents [23, 28-37], and may thus contribute to developmental checkpoints. We therefore took advantage in this work of the strong genetically tractability at the organismal level of *Drosophila melanogaster* to investigate the mechanisms that may allow macrophages to regulate steroid hormone production and developmental growth.

## Results

### Starvation and macrophage-deficiency delay ecdysone biosynthesis and metamorphosis in Drosophila larvae

*Drosophila* larvae lacking either nutrients or hemolectin (hml)-expressing macrophages display a developmental delay in the onset of metamorphosis, also known as the larva-to-pupa transition [27, 38, 39]. Mid-L3 larvae can be starved for up to 8 hrs (see methods, **Fig. S1A**) and survive, but their pupation was delayed by ∼8 hours in comparison to controls fed on standard molasses medium (**Fig. 1A**). Delayed pupation of starved larvae is due to delayed ecdysone biosynthesis by the prothoracic gland [38, 39] (**Fig. S1B**). Accordingly, at the time of the normal onset of pupation in our lab conditions (104 hours A.E.L,) transcription of the ecdysone target *eip75b* (*e75b*) [40] (**Fig. 1B)** and ecdysone levels (**Fig. 1C**) were reduced by ∼90% in comparison to control. Macrophage-less larvae [23, 41, 42] (**Fig. 1C**) also presented with an ∼8 hour delayed pupation, despite control feeding conditions (**Fig. 1D, E**). A time course analysis indicated that transcription of the ecdysone target *eip75b* in these conditions is delayed by ∼8 hours in comparison to controls (**Fig. 1F**), and ecdysone levels are reduced by ∼90% at 104 hours A.E.L. (**Fig. 1G**). Transcription of the ecdysone biosynthetic enzymes [43] *neverland* (*nvd*), *phantom* (*phm*), and *shadow* (*sad*) was also delayed by ∼8 hours (**Fig. 1H)** as observed in starved larvae (**Fig. S1B**). These data suggested that either starvation or macrophage deficiency increases the length of the L3 larval (feeding) stage before pupation by delaying the biosynthesis of ecdysone. Macrophages can act as nutrient sensors [18], and we therefore considered the hypothesis that they may regulate ecdysone biosynthesis to optimize the timing of pupation according to the nutritional status of the larvae (**Fig. 1I**). To test this hypothesis, we performed genetic screens to identify the putative macrophage secreted growth factor(s) or cytokines that could directly or indirectly control developmental timing via ecdysone synthesis, along with the putative sensor(s) for larval nutritional status that could regulate the production of the secreted factor(s) by macrophages.

**Figure 1.**
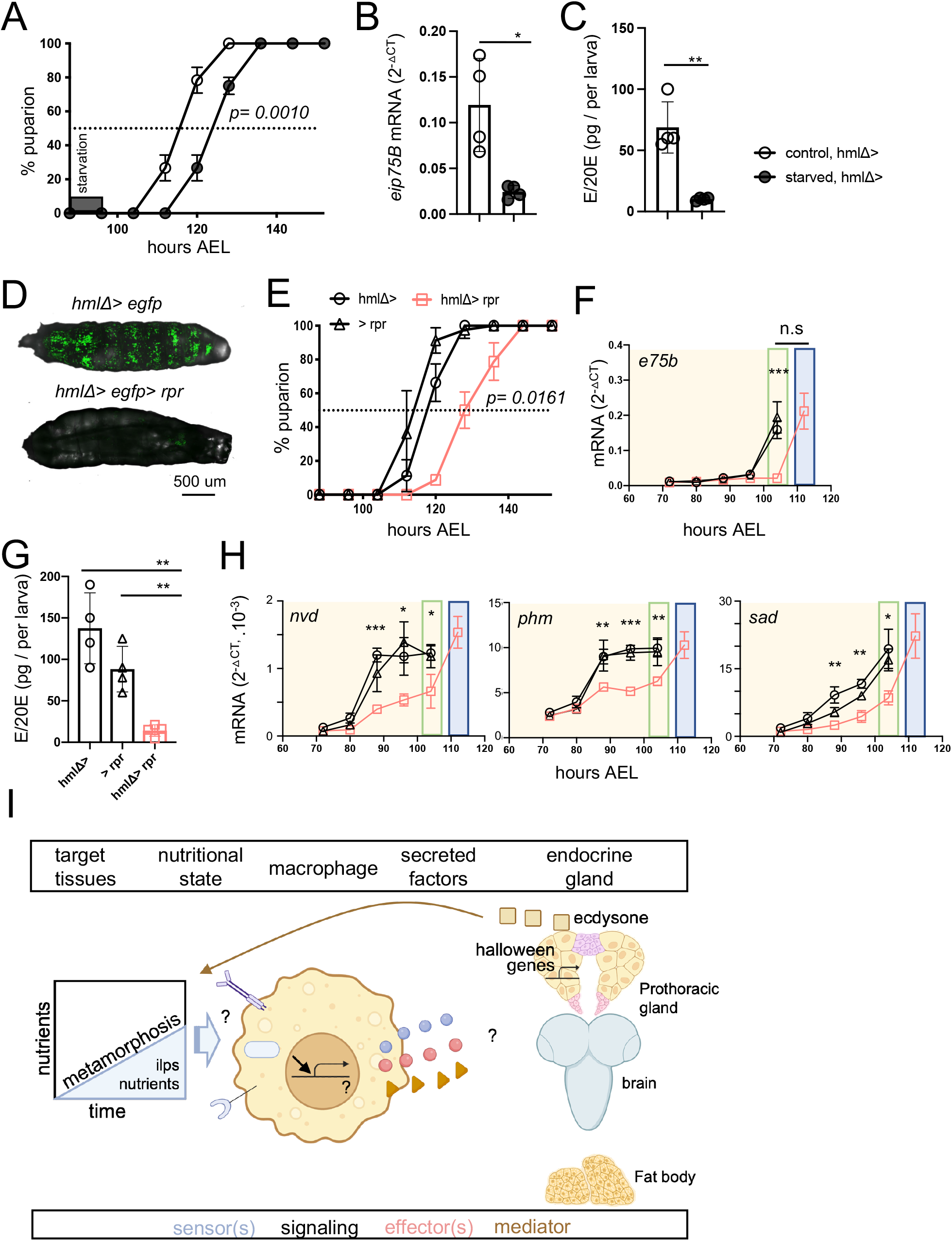
Nutrient deprivation and macrophages-less larvae display a developmental delay from ecdysone synthesis inhibition. (A) Developmental timing in larvae under starvation treatment for 8 hours at 88 hours AEL in the indicate genotypes. (n=3, 20 larvae per genotype in 3 independent crosses, 60 larvae in total per condition). (B) Transcriptional levels of the ecdysone signaling target gene *eip75b* in starved flies. mRNA levels normalized to *rp49* at 104 hours AEL. n=4. Sample were pooled by 5 larvae per condition and genotype. (C) Total ecdysone levels in whole larvae under starvation treatment for 8 hours at 88 hours AEL in the indicate genotypes, quantified by ELISA. n=4, each sample pooled of 10 larvae. (D) Confocal imaging of larvae with GFP fluorescent macrophages and larvae depleted for macrophages. (E) Developmental timing in larvae depleted of macrophages, by overexpression of the reaper pro-apoptotic gene. (n=3, 20 larvae per genotype in 3 independent crosses, 60 larvae in total per condition). (F) mRNA levels of the ecdysone signaling target gene *eip75b* in macrophage-less larvae. mRNA levels are normalized to *rp49* at different time points in larva lacking macrophages. n=3, each sample pooled of 5 larvae. (G) Ecdysone levels in macrophage-less whole larvae, quantified by ELISA. n=4, each sample pooled of 10 larvae. (H) mRNA levels of the Halloween genes *nvd, phm*, and *sad* in macrophage-less larvae. mRNA levels are normalized to *rp49* at different time points in larvae lacking macrophages. n=3, each sample 5 pooled larvae. Green boxes represent the normal developmental timing in control conditions and blue boxes represent the delay in developmental timing in experimental conditions. (I) Working model of ecdysone synthesis regulation by nutrient-sensing receptors and secreted factors present in macrophages during development. Statistical analysis: two-tailed unpaired *t-*test, values represent the mean ± SD. *p < 0.05, **p < 0.01, ***p < 0.001, n.s. not significant.

### Production of the PDGF-family member pvf2 by macrophages controls the timing of ecdysone production and metamorphosis

*Drosophila* macrophages express nutritional sensors, such as nuclear receptors and the insulin receptor, and are a source of secreted factors including cytokines and growth factors that can mediate adaptive responses of local or distant tissues in response to external or internal cues [23, 24, 44]. We thus designed a targeted knock-down screen for genes encoding secreted factors and nutrient-sensing receptors that are expressed in larval macrophages before the onset of pupation (96 hours AEL) [45] (**Fig. S2, S3**), testing their role in the timing of pupation and ecdysone production. We found that selective knock-down of the PDGF/VEGF conserved secreted factor *pvf2* in macrophages, using 3 independent RNAi transgenes and 3 hemocyte drivers recapitulated the ∼8-hour delay in pupation (**Fig. 2A-C & Fig.S4A-C**). Pupation was also delayed in the *pvf2*^*c06947*^ loss-of-function mutant (*pvf2* ^*-/-*^) (**Fig. 2B, Fig.S4D)** [46], while knock-down of *pvf2* in other tissues such as the fat body, prothoracic gland, trachea system and glia cells did not affect pupation (**Fig. 2B, Fig.S4E**). In addition, we found that *pvf2* transcript levels were selectively low in larvae deficient in macrophages (**Fig. 2D)**, and in macrophages from starved larvae (**Fig. 2E, Fig.S4F)**. Furthermore, knock-down of *pvf2* in macrophages delayed transcription of the ecdysone target *eip75b* (**Fig. 2F**), and reduced ecdysone production at 104 AEL (**Fig. 2G**), as observed in macrophage-less and starved larvae. Finally, we found that expression of a *pvf2* transgene in macrophages fully rescued the developmental delay and ecdysone production in starved larvae (**Fig. 2H, I**) as well as in the *pvf2* ^*-/-*^ mutant (**Fig. 2J, K**), although it did not trigger early pupation before 104 AEL (**Fig. 2H, J**). These data altogether indicate that transcriptionally regulated production of Pvf2 by macrophages controls an 8-hour window in the timing of ecdysone production and metamorphosis, based on whether the larvae has eaten. This suggests that production of Pvf2 by macrophages is a nutritional status checkpoint for the developmental timing of metamorphosis.

**Figure 2.**
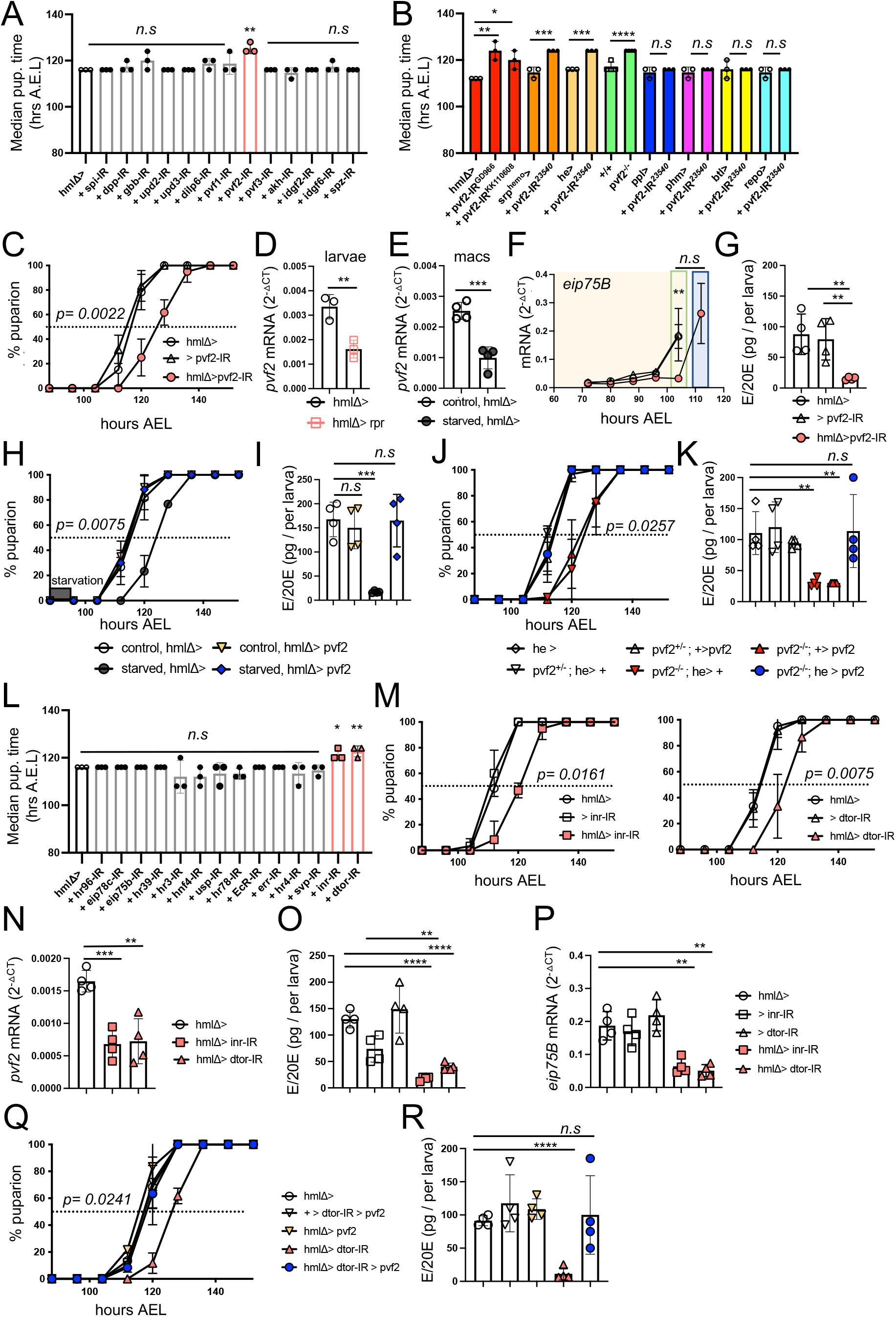
A genetic screen in macrophages identifies a *inr/dtor/pvf2* cassette that controls pupation and ecdysone synthesis under nutritional availability. (A) Median time of puparium formation (hrs AEL) of the secreted factors RNAi screen in macrophages using the macrophage-specific *hmlΔ-gal4* driver and RNAi lines from BDSC. (B) Median time of puparium formation in *pvf2* knock down for two independent RNAi lines (from VDRC) using the macrophage-specific *hmlΔ-gal4* driver, *pvf2* knock down using independent macrophage-specific driver (*srp*^*hemo*^*-gal4*, and *he-gal4*), *pvf2*^*c06947*^ null mutant, and *pvf2* knock down using specific drivers for fat body (*ppl-gal4*), prothoracic gland (*phm-gal4*), trachea (*btl-gal4*), and glia cells (*repo-gal4*) (n=3, 20 larvae per genotype in 3 independent crosses, 60 larvae in total per condition). (C) knock-down of *pvf2* specifically in macrophages induces developmental delay (n=3, 20 larvae per genotype in 3 independent crosses, 60 larvae in total per condition). (D) *pvf2* mRNA levels at 96 hours after egg laying (AEL) of macrophage-less whole larvae (5 larvae per genotype). (E) *pvf2* mRNA levels at 96 hours after egg laying (AEL) of 20000 GFP^+^ macrophages sorted in starved larvae at 96 hours AEL. mRNA levels are normalized to *rp49*. n=3-4. (F) Transcriptional levels of the ecdysone signaling target gene *eip75b* in *pvf2* knock down in macrophage larvae. mRNA levels normalized to *rp49*. n=3. Sample were pooled by 5 larvae per condition and genotype. (G) Ecdysone levels in *pvf2* knock down in macrophage larvae, quantified by ELISA. n=4, each sample pooled of 10 larvae. (H) Developmental timing rescue by overexpression of *pvf2* in flies under starvation treatment for 8 hours at 88 hours AEL in the indicate genotypes. (n=3, 20 larvae per genotype in 3 independent crosses, 60 larvae in total per condition). (I) Ecdysone levels in starved larvae rescued by overexpression of *pvf2*, quantified by ELISA. n=4, each sample pooled of 10 larvae. (J) Genetic rescue of developmental delay in *pvf2*^*c0694*^ loss-of-function mutant by overexpression of *pvf2*, using the macrophage-specific hemmese-gal4 line. (K) Ecdysone total levels rescue in *pvf2*^*c0694*^ loss-of-function mutant by overexpression of *pvf2*in whole larvae, quantified by ELISA. n=4, each sample pooled of 10 larvae of the genotypes indicated. (L) Median time of puparium formation (hrs AEL) of the nutrient-sensing receptors RNAi screen in macrophages using the macrophages-specific *hmlΔ-gal4* driver and RNAi lines from BDSC. (M) knock-down of *inr* and *dtor* specifically in macrophages induces developmental delay (n=3, 20 larvae per genotype in 3 independent crosses, 60 larvae in total per condition). (N) mRNA levels of *pvf2* secreted factor from sorted GFP^+^ macrophages in the indicate genotypes at 96 hours AEL. mRNA levels are normalized to *rp49*. n= 4, each sample pooled of 20000 GFP^+^ macrophages. (O) Ecdysone total levels in whole larvae in the indicate genotype at 104 hours AEL, quantified by ELISA. n=4, each sample pooled of 10 larvae. (P) Transcriptional levels of target gene for ecdysone signaling *eip75b* in flies of the indicate genotypes. mRNA levels normalized to *rp49* at 104 hours AEL. n=4. (Q) Epistatic rescue of developmental timing delay in *dtor*-lacking macrophages mutant by overexpression of *pvf2*. (R) Ecdysone total levels in whole larvae in the indicate genotypes, quantified by ELISA. n=4, each sample a pool of 10 larvae of the indicated genotypes. Data were analyzed by two-tailed unpaired *t-*test and values represent the mean ± SD. *p < 0.05, **p < 0.01, ***p < 0.001, ****p < 0.0001, n.s. not significant.

### Insulin/TOR signaling is required for of Pvf2 production by macrophages

While testing the effect of known molecular sensors of nutritional status, we found that knock-down in larval macrophages of the *insulin-like receptor* (*inr*) [47] and *target of rapamycin* (*dtor)* [48](**Fig.S5A)**, but not of the ecdysone receptor itself, also recapitulated the developmental delay observed in starved, macrophage-less, and macrophage-*pvf2* deficient larvae (**Fig. 2L,M & Fig.S3**). The Insulin/TOR pathway is a conserved sensor of nutritional stress in metazoans, controlling cell and organism growth [49-51] and in particular the timing of differentiation [10]. The array of nutrition-related signaling inputs to this pathway has been defined [52], but the growth-regulatory outputs are less clear. Overexpression of a dominant-negative *inr* (*uas-inr*^*DN*^) in macrophages or a second independent *dtor* RNAi line recapitulated the pupation delay (**Fig. S5B,C**). In addition, macrophage *inr* and *dtor* knock-down both specifically decreased *pvf2* expression (**Fig. 2N, Fig.S5D)**, ecdysone production (**Fig. 2O)**, and transcription of *eip75b* (**Fig. 2P)**. Moreover, we found that the epistatic rescue of *dtor* knock-down larvae by expression of a *pvf2* transgene in macrophages restored developmental timing (**Fig. 2Q**) as well as ecdysone production (**Fig. 2R**). Of note, InR can regulate metabolism and growth via 2 main downstream pathways, activation of Tor and inhibition of FoxO [47], but *FoxO* overexpression did not affect developmental timing (**Fig.S5E)**. These results indicate that the Insulin Receptor/TOR pathway in macrophages regulates ecdysone production and the developmental timing of pupation via the production of Pvf2.

### Macrophage-derived Pvf2 regulates ecdysone biosynthesis via its receptor Pvr in the prothoracic gland

The above data altogether identify an *inr/dtor/pvf2* macrophage genetic cassette that acts as a nutritional status checkpoint that can delay ecdysone production. Ecdysone is mainly produced by the prothoracic gland, an endocrine gland in the *Drosophila* larva [8]. As noted above, ecdysone production in the prothoracic gland is under the control of the *Drosophila* GnRH superfamily hormone Corazonin (Crz)-producing neurons, via the prothoracicotropic hormone (PTTH) produced by the PTTH neurons, and of dILPs produced by Insulin producing neurons [4, 5, 7-9]. However, TK receptors including Pvr can also regulate developmental timing [53]. Pvr is the only known Pvf receptor in *Drosophila* [54], and is expressed ubiquitously by macrophages, neurons, and fat body cells [55, 56]. Thus, we next investigated whether macrophage Pvf2 may control ecdysone synthesis in the prothoracic gland directly or indirectly through its effect on macrophages themselves, neurons, or fat body cells.

A time course analysis of Pvr expression in the prothoracic gland using an anti-Pvr antibody indicated that Pvr is first detected at 88 hours A.E.L and stays detectable until the onset of maturation at 104 hours A.E.L. (**Fig. 3A**). Consistently, we found that transcription of *pvr* in dissected brain/ring gland tissue was upregulated between 80-and 88-hours A.E.L to reach its highest level at 104 hours A.E.L (**Fig. 3B**). Furthermore knock-down of *pvr* in the prothoracic gland recapitulated the 8-hour developmental delay observed in starved, macrophage-less, and *pvf2-, inr-*, and *dtor-*knock-down models above (**Fig. 3C, Fig.S6A**). The hypomorphic *pvr*^*01*^ mutant [54], recapitulated the developmental delay (**Fig. S6B**). Knock-down of *pvr* in fat body, neurons, or macrophages, did not affect developmental timing (**Fig. 3D, Fig. S6C**). In contrast, knock-down of *pvr* in the prothoracic gland did decrease ecdysone synthesis at 104 hrs AEL (**Fig. 3E)** and delayed transcription of the target gene *eip75b* by 8 hours (**Fig. 3F)**. As observed in the starved and macrophage-less *inr* and *dtor* models (see **Fig.1G, Fig. S1B, Fig. S6C**) we found that knock-down of *pvf2* in macrophages and knock-down of *pvr* in the prothoracic gland delayed transcription of the ecdysone biosynthetic pathway genes *nvd, phm*, and *sad* by 8 hours (**Fig. 3G)**. Finally, *pvr* overexpression in the prothoracic gland [53] fully rescued, and even accelerated, developmental timing and ecdysone production in the *pvf2* ^*-/-*^ mutant (**Fig. 3H,I, Fig. S6E,H**). Of note, the latter contrasts with *pvf2* overexpression which did not accelerate developmental timing (**Fig. 2H, J, Fig.S6I**). We also examined a possible redundancy between the role of Pvf2 and its paralogs Pvf1 and Pvf3, as they share the same receptor Pvr [54]. However, starvation and *inr-* and *dtor-*knock-down did not alter *pvf1* and *pvf3* transcript levels (**Fig.S4F, Fig.S5C**). In addition, knock-down of *pvf1* and *pvf3* did not affect developmental timing or ecdysone synthesis (**Fig.2A, Fig.S2D, Fig.S7A-E**). These data altogether indicate that InR/dTor signaling in macrophages is required for Pvf2 production, which is in turn required to trigger pupation at 104 AEL, by acting on its receptor Pvr, expressed in the prothoracic gland at the onset of pupation to initiate ecdysone synthesis. However, these data also reveal that, in the absence of the Pvr-dependent signaling axis, which we describe here, ecdysone biosynthesis and pupation are initiated 8 hours later in a Pvr-independent manner.

**Figure 3.**
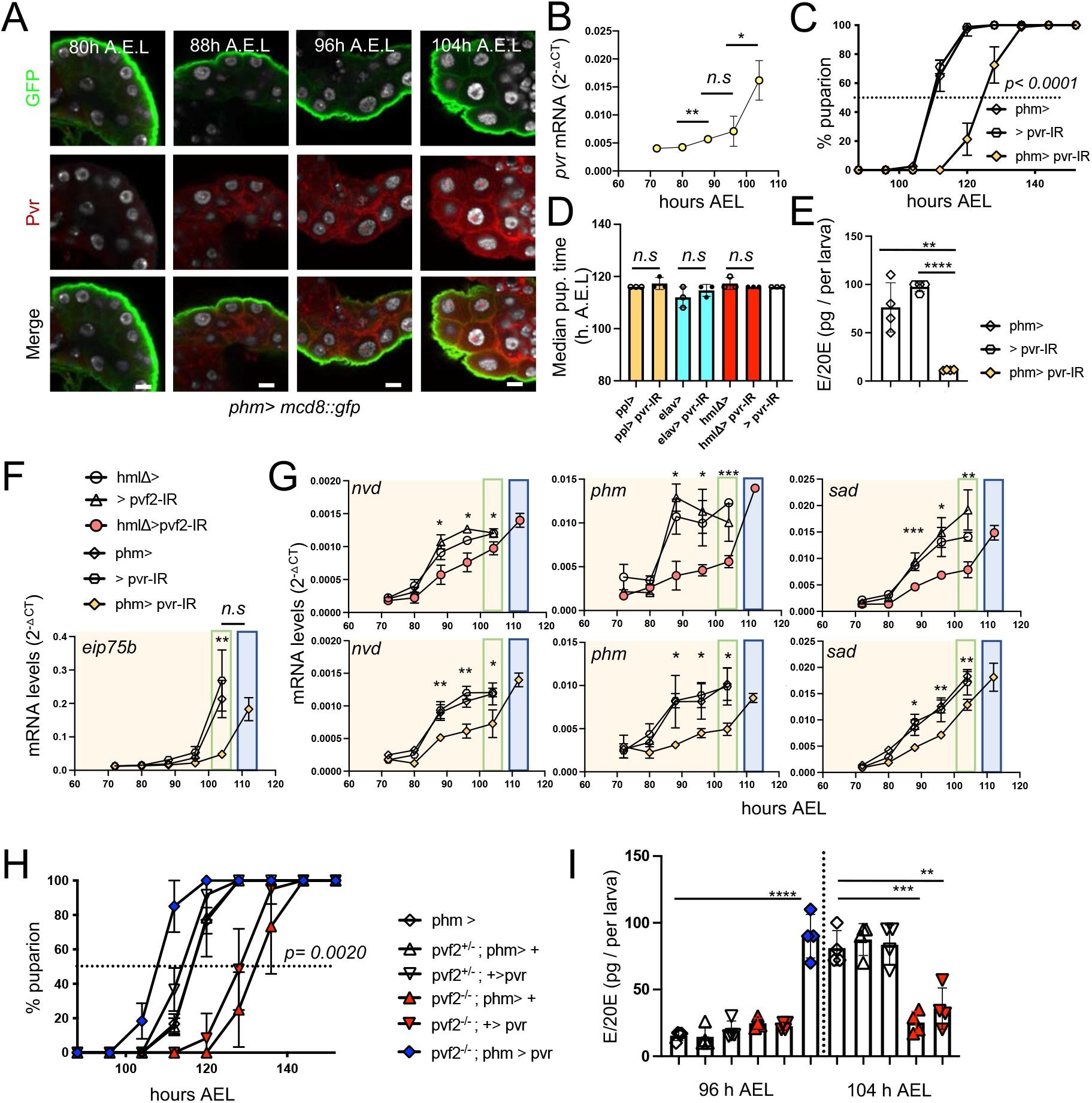
Macrophage-derived Pvf2 acts through its receptor Pvr in the prothoracic gland to control ecdysone synthesis. (A) Pvr temporal expression pattern in prothoracic gland at 80, 88, 96, and 104 hours AEL. Prothoracic gland cellular membranes (Green) begin to express Pvr (Red) at 88 hours AEL, and continue to until the onset of pupariation at 104 hours AEL. Scale bars are 20 um. n= 4-7 for each time point. (This is an antibody stain? Or in situ? I would clarify what was done) (B) *pvr* temporal transcriptional levels in third instar-larvae at 72, 80, 88, 96 and 104 hours AEL in whole larva control (w^1118^). mRNA levels are normalized to *rp49*. (C) knock-down of *pvr* specifically in the prothoracic gland induces developmental delay (n=3, 20 larvae per genotype in 3 independent crosses, 60 larvae in total per condition). (D) Median time of puparium formation in *pvr* knock-down specifically in fat body (*ppl-gal4*), and neurons (*elav-gal4*), and macrophages (*hmlΔ-gal4)* (n=3, 20 larvae per genotype in 3 independent crosses, 60 larvae in total per condition). (E) Total ecdysone levels in whole larvae in the indicate genotype at 104 hours AEL, quantified by ELISA. n=4, each sample pooled of 10 larvae. (F) Ecdysone target gene *e75b* mRNA levels normalized to *rp49* at different time points in larva lacking *pvr* in prothoracic gland. n=3, each sample a pool of 5 whole larvae at each time point with indicated genotype. (G) mRNA levels of the Halloween genes *nvd, phm*, and *sad* in larvae *with pvf2* knocked down in macrophages or *pvr* knocked down in the prothoracic gland. mRNA levels are normalized to *rp49* at different time points in larva lacking macrophages. n=3, each sample a pool of 5 larvae. Green boxes represent the normal developmental timing in control conditions and blue boxes represent the delay in developmental timing in experimental conditions. (H) Epistatic rescue of developmental timing delay in *pvf2*^*c0694*^ loss-of-function mutant by overexpression of *pvr*, using the prothoracic gland-specific line *phantom-gal4*. (I) Total ecdysone levels in whole larvae in the indicated genotypes, quantified by ELISA. n=4, each sample a pool of 10 larvae of the genotypes indicated. Data were analyzed by a two-tailed unpaired *t-*test and values represent the mean ± SD. *p < 0.05, **p < 0.01, ***p < 0.001, ****p < 0.0001. n.s (not significant).

### Pvf2 biological activity is regulated by transcription rather than by macrophages numbers

Pvf2 promotes proliferation and survival of macrophages via Pvr in a cell autonomous manner [46, 54, 57], which represents a possible confounding factor in the above analysis. Macrophages are indeed reduced by ∼30% in *pvf2* macrophage knock down larvae (**Fig. 4A**). Thus, manipulating this axis could alter signal production by macrophages by altering their quantity. However, *pvr* knock down in hemocytes, which also reduces macrophage numbers by ∼30% (**Fig. 4A)**, does not decrease *pvf2* production by macrophages nor does it delay pupation, in comparison to controls (**Fig. 4B**). Macrophages numbers are also reduced by ∼30% in starved larvae as well as macrophage-*inr* and *dtor* knock-down larvae (**Fig. 4A**). However, RT-qPCR quantification showed a 60 to 80% decrease, on a per cell basis, of *pvf2* transcripts in FACS-sorted macrophages from starved, *inr, dtor*, and *pvf2* knock-down larvae (**Fig. 4B**). This indicates that Pvf/Pvr signaling in macrophages regulates their numbers but does not control *pvf2* expression, while starvation, InR, and dTor do. Therefore, starvation, as well as *inr, dtor*, and *pvf2* knock-down moderately decrease macrophage numbers, which may contribute to the overall decrease in *pvf2* production observed in the starved and mutant larvae but is unlikely to account for the biological activity of Pvf2 in the control of ecdysone synthesis and metamorphosis. Nevertheless, the production of Pvf2 could be restricted to a subset of macrophages whose survival or proliferation is selectively affected by starvation and deficient InR/dTor, and Pvr signaling. Analysis of *inr, dtor, pvr* and *pvf2* expression in a recent *Drosophila* larva single cell RNAseq dataset [58] showed that *inr, dtor*, and *pvr* expression is widely distributed and is not restricted to any potential subset (**Fig. 4C**). Interestingly *pvf2* expression itself, although distributed throughout many hemocyte clusters, appears more abundant on a per cell basis in clusters of activated hemocytes (LM1, PM5, PM3 and PM8), consistent with the idea that a threshold of hemocyte activation is associated with *pvf2* transcription. Altogether, our data indicate that InR and dTor signaling in macrophage license ecdysone synthesis via the transcriptional regulation of *pvf2*, and to a lesser extent by the regulation of hemocyte numbers themselves. These data are consistent with the hypothesis that the expression or lack of expression of *pvf2* by macrophages acts as a developmental checkpoint that licenses Pvr-dependent pupation if the larval nutritional status, as monitored by InR and dTor, is adequate.

**Figure 4.**
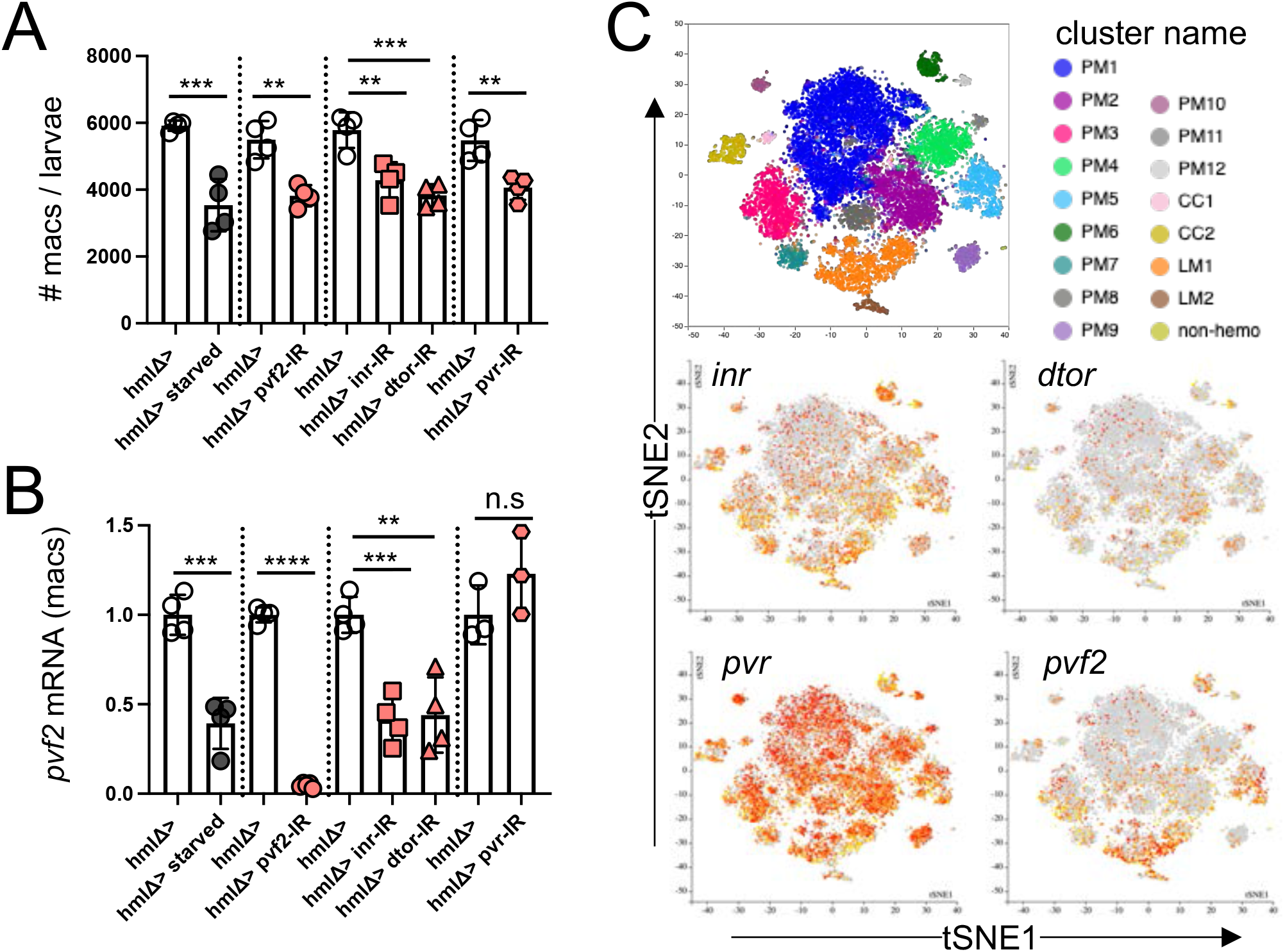
The macrophage *Inr/dtor/pvf2* cassette control macrophage numbers and *pvf2* expression. (A) Quantification of macrophage numbers in total larvae at 96 hours AEL by FACS-sorted GFP^+^ cells in the indicated genotypes. Dots represents results from groups of 5 larves, n=4 independent experiments. (B) Dots represent normalized *pvf2* mRNA levels at 96 hours after egg laying (AEL) in 20,000 GFP^+^ macrophages sorted from 15 to 20 larvae of the indicated genotypes. n=3 to 4 independent experiments. mRNA levels are normalized to *rp49*. (C) t-SNE plots from single-cell RNA sequencing expression of *inr, dtor, pvr* and *pvf2* in wandering larva hemocytes (from Ref. 58, see Methods). Data were analyzed by a two-tailed unpaired *t-*test and values represent the mean ± SD. **p < 0.01, ***p < 0.001, ****p < 0.0001. n.s (not significant).

### The macrophage Inr/dtor/pvf2 cassette control the size of adult flies

Delays in critical transitions such as pupation have systemic effects on organismal and tissue growth [4, 13, 39, 53]. In particular delayed pupation in response to suboptimal food supplies is believed to minimize the effect of nutrient deficiency on the size and fitness of the adult by extending the growth period [51]. We thus investigated the consequences of the macrophage- and pvf2/pvr-dependent developmental delay for the size of pupae and adult *Drosophila*. We found that abrogating the pupation delay in starved larvae by a *pvf2* transgene further reduced the size of the pupae (**Fig.5A**). Conversely, enforcing a Pvr-dependent developmental delay by knock-down or expression of dominant negative forms of *inr, dtor* and knock-down of *pvf2* in macrophages using several macrophage specific drivers and *pvf2* RNAi lines and in pvf2^-/-^mutants, always resulted in larger pupae under *ad libitum* feeding condition (**Fig.5B,C,D)**. In addition, the genetic rescue of pvf2^-/-^ mutants, and the epistatic rescue of *dtor* knock-down larvae by overexpression of a *pvf2* transgene both restored wild type size (**Fig.5D**). In contrast, knock-down of *pvf2* in glia, prothoracic gland, trachea, and fat body had no effect on pupal size (**Fig.5E)**. Knock-down of *pvf1* and *pvf3* in macrophages had no effect on pupal size either (**Fig.S7F**). Finally, the knock-down of *inr, dtor*, and *pvf2* in macrophages also generated larger adult flies, both female and male, as measured by wing size (**Fig.5F**). To further confirm that the effect of the *inr/dtor/pvf2* cassette on pupal and adult size is mediated by Pvr signaling in the prothoracic gland, we examined the role of *pvr* knock down, loss of function, or gain of function. Knock-down of *pvr* in the prothoracic gland -but not in macrophages-increased the size of pupae (**Fig.5G)**. In addition, the pvr hypomorph mutant phenocopied the knock-down of *pvr* in the prothoracic gland (**Fig.5G**). Knock-down of *pv*r in the prothoracic gland also increased the sizes of adult female and male flies (**Fig.5H**). In contrast, overexpression of *pvr* in the prothoracic gland, which shortens the pupation time (see Fig. 3G and Fig. S6E), decreased the size of pupae (**Fig.5I)** and adult flies (**Fig.5H**). Finally, the size increase observed in null *pvf* ^*-/-*^ mutants was epistatically rescued by *pvr* overexpression in the prothoracic gland (**Fig.5I**). These data altogether demonstrate that control of the Pvr-dependent ecdysone synthesis and pupation delays by the macrophage *inr/dtor/pvf2* cassette regulates the size of pupae and adult flies from both sexes (**Fig.5J**).

**Figure 5.**
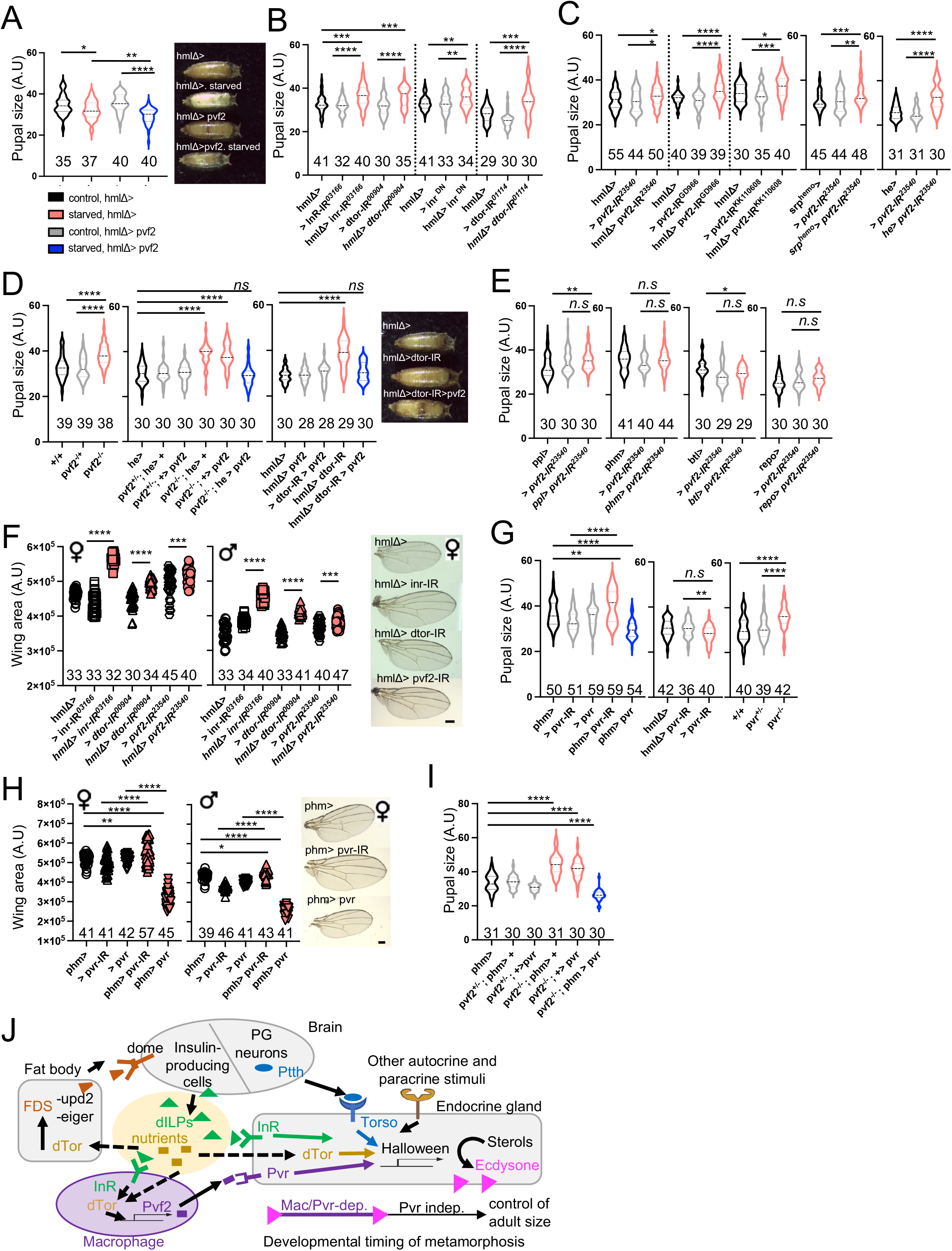
The macrophage *Inr/dtor/pvf2* cassette control adult size by Pvr in prothoracic gland. (A) Final pupal size of starved larvae in the indicated genotypes. (B) Final pupal size of *inr* and *dtor* knock-down in macrophages. (C) Final pupal size of *pvf2* knock-down using different RNAi lines and macrophage drivers. (D) Final pupal size in *pvf2* ^*c06947*^ loss-of-function flies, genetic rescue of pupal size in *pvf2*^*c0694*^ loss-of-function mutant by overexpression of pvf2, and epistatic rescue of pupal size in *dtor*-lacking macrophages mutant by overexpression of *pvf2*. (E) Final pupal size upon *pvf2* knock-down in fat body (*ppl-gal4)*, prothoracic gland (*phm-gal4)*, trachea (*btl-gal4)*, and glia cells *(repo-gal4)*. (F) Wing area of female and male virgin flies of *inr, dtor*, and *pvf2* knock-down in macrophage. (G) Final pupal size upon *pvr* knock-down or *pvr* overexpression in prothoracic gland, knock-down of *pvr* in macrophages, and in *pvr*^*01*^ hypomorph mutant. (H) Wing area of female and male virgin flies upon *pvr* knock-down or *pvr* overexpression in prothoracic gland. (I) Epistatic rescue of pupal size in *pvf2*^*c0694*^ loss-of-function mutant by overexpression of *pvr* in prothoracic gland. Numbers along the x axis of the graphs indicate the number of flies analysed. Data were analyzed by a two-tailed unpaired *t-*test and values represent the mean ± SD. *p < 0.05, **p < 0.01, ***p < 0.001, ****p < 0.0001. n.s (not significant). (J) **Working model** for the role of the *Inr/dtor/pvf2* cassette in macrophages (purple colors) as a nutritional status checkpoint in the context of current knowledge of the timing of metamorphosis by the brain, the fat body, nutritional status and steroid hormone biosynthesis in the prothoracic gland. Purple arrows indicate the control of a Pvr-dependent stage of ecdysone biosynthesis under the control of macrophages.

## Discussion

### The macrophage inr/dtor/pvf2 cassette is a developmental timing checkpoint which increases fitness

We report here that validation of the nutritional status of the larvae by the macrophage *inr/dtor/pvf2* genetic cassette is a checkpoint that allows Pvr-dependent ecdysone synthesis in the prothoracic gland and the onset of metamorphosis to proceed. If this nutritional status is not met, here in the case of a starvation event, low Pvf2 production delays ecdysone synthesis and opens a time-window of an extra ∼8 hours for preparing pupation, after which Pvr-independent ecdysone synthesis and pupation occur (**Fig.5J**). The physiological significance of this time-window controlled by macrophages and Pvr expression in the prothoracic gland for the fitness of the host is best illustrated by its consequence on the fitness, here the size, of pupae and adult flies, as opening the window allows the formation of larger flies and can rescue the effect of a starvation episode at the larval stage [38, 51].

These findings support and extend the general concept that macrophages can act as metabolic sensors which integrate environmental signals to contribute to organismal homeostasis [18, 22]. More precisely, the results from this study shows that macrophages represent a step in a complex and highly regulated inter-organ communication process that allows for fine-tuning of developmental timing. The role of macrophages appears to be to delay the timing of the developmental program when the nutritional status of the larvae is poor, as assessed by InR/dTor signaling, but early/increased *pvf2* expression does not initiate premature Pvr-mediated pupation. In contrast, the timing of Pvr-mediated pupation is determined by expression of Pvr in the prothoracic gland, which is upregulated just before the onset of normal pupation, while *pvr* early/overexpression induces early pupation and strongly decreases the size of the adult flies. Moreover, in the absence of Pvf2 or Pvr, pupation will eventually proceed, in a Pvr-independent manner, at the cost of a reduced size of the flies.

### Macrophage assess nutritional status via InR/Tor signaling

Our results suggest that macrophages sense the nutritional status of the larvae via Insulin/TOR signaling. This is consistent with seminal studies which have shown that the Insulin/TOR signaling network transduces nutritional signals that regulate cell growth and developmental timing at the organismal level [10, 13, 59, 60]. DILPs synthesis and secretion by insulin-producing cells – like insulin in mammals- [47, 61] is stimulated directly and indirectly by fat body secreted factors such as Unpaired 2, Eiger, and Stunted which are decreased in starvation conditions [62-65] (**Fig.5J**). DILPs circulate in the hemolymph and are thus available to macrophages. In addition, dTor -as well as the mammalian TOR complex-is activated by Insulin receptor signaling and also directly by nutrients, specifically amino-acids [48]. The insulin receptor also signals independently of Tor by inhibiting Foxo [47], but our results indicate that FOXO overexpression does not affect the timing of pupation (see **Fig.S5D**), while dTOR is both necessary and sufficient for Pvf2 production by macrophages and for pupation. We did not observe an effect of the knock down of individual nuclear receptors in macrophages (see **Fig.S3**), suggesting they do not regulate Pvf2 production. Therefore, it is likely that Insulin/TOR signaling in hemocytes is activated, at least in part, by the availability of DILPs and amino acids; nevertheless, it is very possible that macrophages integrate additional, yet unidentified signals, to control *pvf2* transcription and steroid hormone biosynthesis.

### Bioavailability of Pvf2 in the prothoracic gland

*Drosophila* Pvr has 3 ligands coded by separate genes, *pvf1, pvf2* and *pvf3*, with partially redundant functions in cell survival, migration, and proliferation during embryonical stages [46, 54, 66, 67]. Nevertheless, our results indicate that only Pvf2 can activate Pvr in the prothoracic gland to regulates ecdysone synthesis and developmental timing under physiological conditions **(Fig.2, and S7)**. Among possible explanations, we show here that *pvf2* expression is 10-to 100-fold higher in larval macrophages as compared to *pvf1* and *pvf3* **(Fig.S2C)**. Nevertheless, production of Pvf1, Pvf2, and Pvf3 by other tissues might have compensated for the deficiency in macrophage Pvf2, which is not observed. It is possible that larval macrophages that produce Pvf2 are in close contact with the prothoracic gland, or that Pvf2 produced by hemocytes circulates in hemolymph and thus access the prothoracic gland more readily than Pvfs produced by other tissues. Finally, we cannot eliminate the possibility of unknown biochemical differences between Pvfs that could affect, for example, their solubility in the hemolymph, of the existence of putative co-receptor(s).

### Macrophages license steroid hormone biosynthesis in the endocrine gland

Steroid hormone production in the endocrine gland of the fly and metamorphosis are under the control of the GnRH superfamily hormone Corazonin (Crz), via the prothoracicotropic hormone (PTTH) produced by the PTTH neurons [4, 7-9]. A significant finding from our study is that the production of Pvf2 by macrophages contributes orthogonally to steroidogenesis by promoting transcription of the Halloween steroid biosynthesis genes when a satisfactory nutritional status is detected (**Fig.5J**). The development and organismal homeostasis of *Drosophila* and mammals are highly conserved [68], and the production of steroid hormones in mammalian puberty is under the control of the hypothalamus gonadotropin-releasing hormone (GnRH) via the anterior pituitary hormones LH, FSH [2, 3, 6], which are similar to the Crz-Ptth axis [9] although LH, FSH are not Ptth orthologs. Although an orthogonal role of macrophages in the regulation of puberty is yet unknown, it is well documented that macrophage-deficient mice present with steroid hormone deficiency and infertility [69-76], and several mouse PVFs orthologs (PDGFs) are also important for steroid production and male and female fertility [77].

## Authors contribution

SJC and FG designed the study and wrote the manuscript. SJC performed all experiments and prepared figure panels.

## Acknowledgements

This work was supported by NIH/NCI P30CA008748 to MSKCC and by NIH/NIAID 1R01AI130345, NIH/NHLBI R01HL138090, Leducq transatlantic network of excellence, Ludwig institute for Cancer research basic immunology grant to F.G. S.JC. is the recipient of a CRI Irvington postdoctoral fellow (CRI award 3440). Authors are indebted to the Bloomington Stock Center at Indiana University (Bloomington, IN) and the Vienna *Drosophila* Resource Center (VDRC), and to Dr. Daria Siekhaus for sharing her RNAseq gene expression analysis of embryonic hemocytes, Dr. Benny Shilo for the kind gift of his rat anti-Pvr antibody, Dr. Bruno Lemaitre for the kind gift of UAS-pvf2 lines, Dr. Sergey Sinenko for for the kind gift of *hmlΔ-gal4;UAS-2xeGFP*, and Dr. Nobert Perrimon for for the kind gift of *srp*^*hemo*^*-gal4* line. Authors are also indebted to the members of the Geissmann lab for helpful discussion and to Drs. Daria Siekhaus, Marco Milan, Maria Dominguez, Chris Glass and Nehemiah Cox for critical reading and review of the manuscript before submission.

## Materials and Methods

### Drosophila husbandry

Plasmatocyte specific reporter line and driver hmlΔ-gal4;UAS-2xeGFP stock is a gift from Sergey Sinenko [1], UAS-pvf2 is a gift from Bruno Lemaitre [2]. srphemo-gal4 is a gift from Nobert Perrimon. spi-TRiP.HMS01120-RNAi, gbb-TRiP.HMS01243-RNAi, dpp-TRiP.JF01090-RNAi, pvf1-TRiP.HMS01958-RNAi, pvf2-TRiP.HMJ23540-RNAi, pvf3-TRiP.HMS01876-RNAi, dilp6-TRiP.HMS00549-RNAi, spz-TRiP.HM05024-RNAi, upd2-TRiP.HMS00948-RNAi, upd3-TRiP.HM05061-RNAi, akh-TRiP.HMS00477-RNAi, idgf2-TRiP.HMC04223-RNAi, idgf6-TRiP.HMC04127-RNAi, pvr-TRiP.HMS01662-RNAi, hr96-TRiP.JF02350, eip78c-TRiP.JF02258, eip75b-TRiP.HMS01530, hr39-TRiP.JF02432, hr3-TRiP.HMC03156, hnf4-TRiP.JF02539, usp-TRiP.HMS01620, hr78-TRiP.JF03424, ecr-TRiP.HMJ22371, err-TRiP.HMC03087, hr4-TRiP.HM05260, svp-TRiP.JF03105, inr-TRiP.HMS03166, elav-gal4, btl-gal4, phm-gal4, ppl-gal4, repo-gal4, he-gal4, UAS-reaper, UAS-inrDN, dtor-TRiP.HMS00904, dtor-TRiP.HMS01114, UAS-pvr, Pvf2c06947, Pvr1, UAS-pvf1.RB, pvf2d02444 lines are from the Bloomington Stock Center at Indiana University (Bloomington, IN). pvf1-KK112211-RNAi, pvf1-GD3432-RNAi, pvf2-KK110608-RNAi, pvf2-GD966-RNAi, pvf3-KK112796-RNAi, and pvf3-GD5238-RNAi are from Vienna *Drosophila* Resource Center (VDRC).

Flies were reared in standard molasses fly food from Archon scientific at 25°C (except when indicated) on a 12:12-hour light:dark cycle.

### Egg-laying and Puparium formation measurements

We crossed 20 females and males and, after 48 hours, the parental flies were transferred to grape agar plates (Nutri-fly grape agar premix from Genesse scientific) with yeast paste and left 3-4 hours for egg deposition at 25°C. The flies were removed, and laid eggs were incubated 48 hours at 25°C. 48 hours later, the second-instar larvae were transferred onto *Drosophila* molasses food (20 larvae per tube) and reared at 26.5°C. We check the transition between larva-to-pupa at 8-hour intervals, with “time 0” designated as 4 hours after the initiation of egg laying and referred to as “after egg laying (A.E.L)”. Median time of pupation was used to analyzes differences between genotypes (50% of larvae that became pupae).

### Pupae volume measurements

20 females and 20 males were crossed and left 24 hours for egg deposition. Parental flies were transferred every 24 hours to fresh tubes, and laid eggs were reared at 25°C. Pupae were collected and photographed with their dorsal side up. Length and width were measured using ImageJ; volume was calculated according to the following formula: v = 4/3π(L/2)(l/2)2 (L, length; l, width) [3]. Leica M80 stereo microscope was used for pupal imagine.

### Adult wing area measurements

20 females and 20 males were crossed and left 24 hours for egg deposition. Parental flies were transferred every 24 hours to fresh tubes, and laid eggs were reared at 25°C. Adults were collected and left wings of each individual were excised and rinsed thoroughly with ethanol and mounted in a glycerol-ethanol solution. Wing areas were measured using ImageJ [4]. Pictures were taken using Axio Lab.A1 Zeiss microscope with N-Achroplan 5x/01.25 objective.

### Whole larva mounting and confocal imaging

Mild-late L3 larvae were rinsed with tap water (no longer than 5 min) dried carefully and anaesthetized using diethyl ether for less than 3.5 min. Larva were positioning within one cover slide and a slide for mounting immobilizing the sample and imagine. We set up the tile scan with the µm thick Z-stack on LSM880 Zeiss microscope with 5x objective, that allows to see the sessile macrophages in the cuticle. Images were acquired with a resolution of 1024×1024.

### Immunohistochemistry in ring-gland tissue

Brain-ring gland complex were dissected out in cold phosphate-buffered saline (PBS) buffer and fixed in 4% paraformaldehyde for 20 minutes [5]. Brain-ring gland complex were stained overnight at 4ºC with chicken anti-gfp (1/500, Life Technologies), and rat anti-pvr [6] (1/500, a gift from Benny Shilo). Secondary antibodies 488 anti-chicken (1/500) and 647 anti-rat (1/500) were purchased from Invitrogen. For DNA and nuclei staining, we used Hoechst (1/1000) for 10 min at room temperature. We set up the tile scan with the µm thick Z-stack on LSM880 Zeiss microscope with 40x oil objective. Images were acquired with a resolution of 1024×1024.

### Macrophages counts

Hemocytes from 5 larvae at 96 hours AEL were bled into 120 µL of FACS buffer (PBS, 0.5% BSA, and 2mMEDTA) containing 1 nm phenylthiourea (PTU, Sigma, St-Louis, MO, USA) to prevent melanization. We used 2 forceps to make an incision from the posterior and pull to the anterior ends of the larvae. We allow larvae to bleed for a few seconds and later we scrape the lateral cuticle, with the careful to avoid the lymph-gland disruption [2]. We counted the macrophages numbers by sorted live GFP positive hemocytes (DAPI negative), using HmlΔ>eGFP, in on an ARIA III (BD bioscience). Flow cytometry plots were obtained with Flow Jo 9.9. (Did you do this count also on the hemocyteless larvae, as it looks in Figure 1 D as if there is residual punctate GFP expression, which may or may not be some hemocytes).

### May-Grunwald Giemsa staining

Analysis of macrophages was done on of sorted cells from GFP positive hemocytes population, using HmlΔ>eGFP. 1000 cells were sorted directly in FBS (Invitrogen) and centrifuged for 10 min at 800 g with low acceleration onto Superfrost slides (Thermo Scientific). After air drying for at least 30 min, the slides were fixed in methanol, air dried for at least 30 min, stained with May-Grunwald (Sigma) solution for 5 to 15 min then Giemsa (Sigma) solution 14% for 15 to 30 min, and rinsed with Sorenson buffer pH 6.8. After air drying, the slides were mounted with Entellan (Merck). Pictures were taken using Axio Lab.A1 Zeiss microscope with N-Achroplan 100x/01.25 objective.

### qRT-PCR on sorted macrophages and whole larvae

For qRT-PCR, 20000 hemocytes were sorted into 600μL of Trizol LS (Life Technologies). RNA extraction was performed following manufacturer’s instructions (miRNeasy Mini Kit Quiagen). RNA concentration was measured with nanodrop2000. cDNA preparation was performed with Invitrogen SuperScript IV First Strand Synthesis System (Thermo Fisher Scientific) as per manufacturer’s instructions. qRT-PCR were done with 5-10 ng cDNA. For whole larvae qRT-PCR were performed tissue homogenization with tissue lyser (Precellys evolution, Bertin technologies) in 2ml tissue homogenizing mixed beads (Bertin technologies, ref: P000918-LYSK0-A) of 5 larvae per sample, and RNA extraction was performed following manufacturer’s instructions (RNeasy Mini Kit Quiagen). For whole larva the same protocol as for the sorted cells and the qRT-PCR was performed on 500ng cDNA. qRT-PCR were performed using Quant studio 6 flex real time PCR from Life Technologies.

qRT-PCR are performed on a Quant Studio 6 Flex using TaqMan Fast Advance Mastermix, and TaqMan probes for rp49 (Dm02151827_g1), dib (Dm01843083_g1), phm (Dm01844265_g1), nvd (Dm03419116_m1), sad (Dm02139323_g1), spo (Dm01840221_s1), eip75b (Dm01793666_m1), idgf2 (Dm01842858_g1), idgf6 (Dm01842553_g1), gbb (Dm01843010_s1), dpp (Dm01842959_m1), akh (Dm01822073_g1), spi (Dm02364865_s1), spz (Dm02151534_g1), spz2 (Dm01832735_g1), spz3 (Dm01809417_m1), spz4 (Dm01810735_g1), spz5 (Dm01835260_m1), spz6 (Dm01834239_g1), dilp6 (Dm01829746_g1), upd2 (Dm01844134_g1), upd3 (Dm01844142_g1), pvf2 (Dm01814370_m1), pvf1 (Dm01814376_m1), pvf3 (Dm01813949_m1), pvr (Dm01803625_m1), hr96 (Dm02151377_m1), eip78c (Dm01798025_g1), hr39 (Dm01811265_g1), hr3 (Dm01846760_m1), hnf4 (Dm01803764_g1), usp (Dm01841709_s1), hr78 (Dm01798077_g1), ecr (Dm01811604_g1), err (Dm01835887_g1), hr4 (Dm01798975_g1), hr38 (Dm01842602_g1), hr83 (Dm02137429_g1), hr51 (Dm01824332_m1), svp (Dm02135706_m1), dsf (Dm01842632_m1), tll (Dm02151861_g1), inr (Dm02136218_g1), dtor (Dm01843300_g1), 4e-bp (Dm01842928_g1).

### Ecdysteroids measurement

For ecdysteroids extraction, 10 whole larvae from each condition (from 96 hours or 104 hours AEL, as indicate in the figure) were collected in 2ml tissue homogenizing mixed beads (Bertin technologies, ref: P000918-LYSK0-A) and frozen in liquid nitrogen and transfer to -80°C. Samples were homogenized using tissue lyser (Precellys evolution, Bertin technologies) in 0.3 ml of methanol, centrifuged at maximum speed for 5 min at room temperature (RT) and the supernatant was transferred to a new eppendorf tube. We repeat this procedure by adding 0.3 ml of methanol to the pellet and mix using vortex and a third round using 0.3 ml of ethanol. The pooled sample after homogenization was 0.9 ml per sample. For quantification, the extracted samples were centrifuged at maximum speed for 5 min (RT) to remove any remaining debris and divided into two tubes to generate technical replicates. The cleared samples were evaporated overnight [7]. The following steps were performed using the 20-hydroxyecdysone ELISA kit instructions (Bertin Pharma #A05120.96 wells). Absorbance was measured at 410 nm with Microplate reader (SpectraMax i3 from Molecular devices).

### Starvation

We followed the egg-laying procedure as explain above. After 48 hours A.E.L, we transferred the larva into petri dishes with *Drosophila* molasses food (20 larvae per dish), and we allowed the larva to develop. At the 88 hours A.E.L, we transferred the larva to empty petri dished with a wet paper in the cover to keep larvae hydrated and avoid desiccation for 8 hours to starve the larvae [8]. After 8 hours, we refeed the larvae transferring then to *Drosophila* molasses food tubes with 5 ml, and we counted puparium formation in 8 hours intervals.

### Single-cell gene expression visualization by dot plots

Dot plots were generated using a user-friendly searchable web-tool (www.flyrnai.org/scRNA/blood/) where genes can be queried and visualized in the different clusters made by single-cell RNA sequencing [9].

### Statistical Analyses

All statistical analyses were performed using GraphPad Prism Software 9.0 with a 95% confidence limit (p < 0.05). An unpaired t test was used for comparisons between two genotypes or time-points.

---

**Figure S1.**
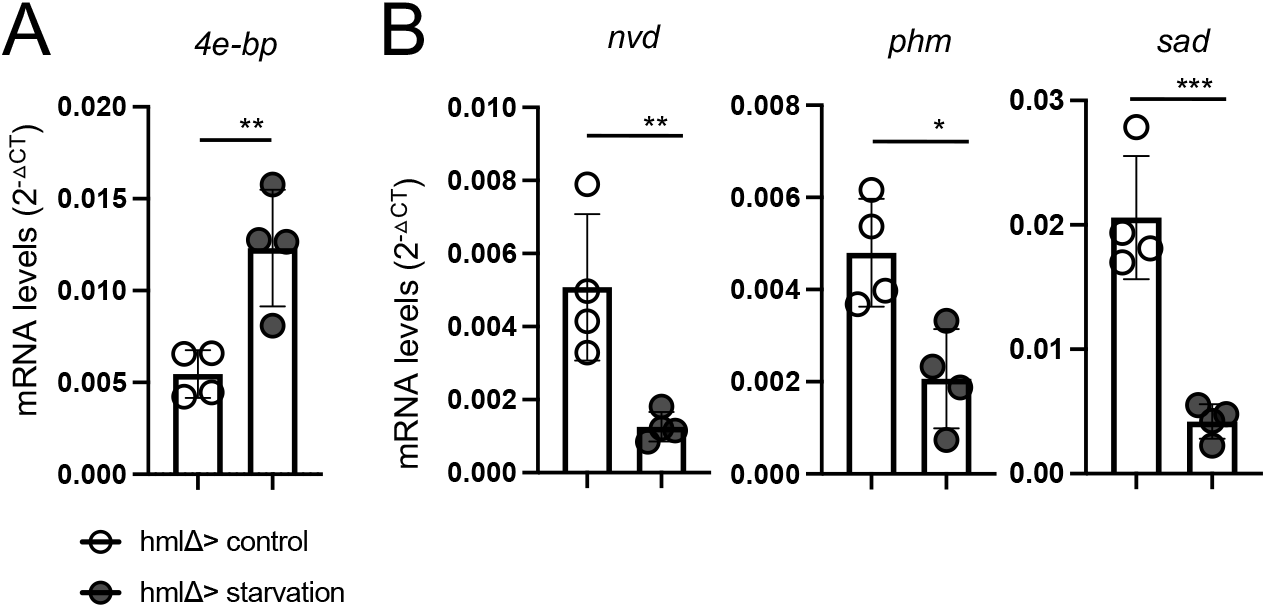
Insulin signaling and ecdysone biosynthesis in starved larvae. (A) Transcriptional levels of the direct target of the growth inhibitor FoxO gene *Thor/4e-bp* in starved flies. mRNA levels normalized to *rp49* at 104 hours AEL. n=4. (B) Transcriptional levels of the ecdysone biosynthesis gene *nvd, phm*, and *sad* in starved larvae. mRNA levels normalized to *rp49* at 104 hours AEL. n=4. Sample were pooled by 5 larvae per condition and genotype. Data were analyzed by a two-tailed unpaired *t-*test and values represent the mean ± SD. *p < 0.05, **p < 0.01, ***p < 0.001, ****p < 0.0001. n.s (not significant).

**Figure S2.**
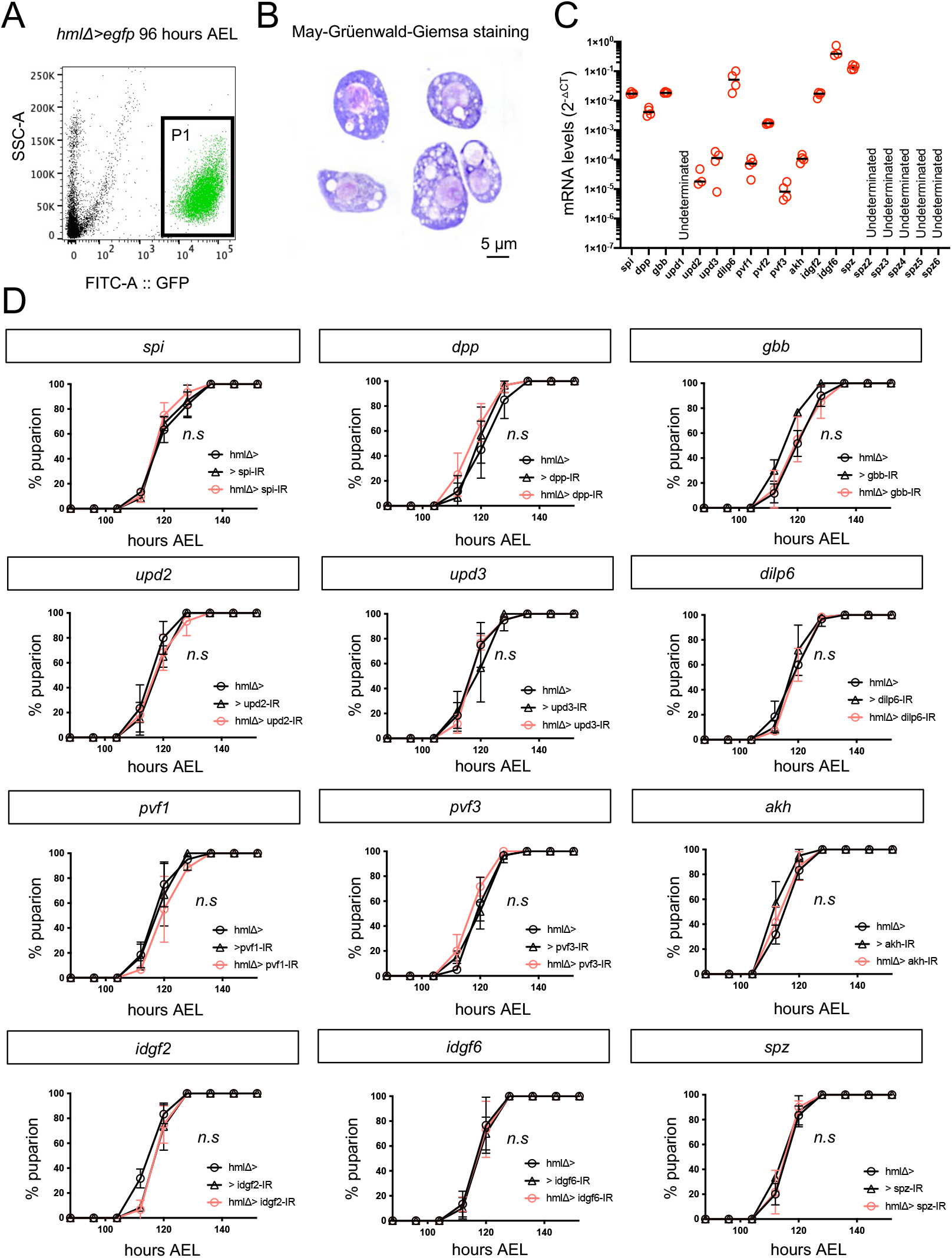
Screen for secreted factors expressed by larval hemocytes. (A) FACS-sorting strategy of GFP^+^ macrophages labeled by the hemolectinΔ-gal4 line. (B) May-Grüenwald-Giemsa staining for GFP^+^ macrophages labeled by the hemolectinΔ-gal4 line. (C) Candidate secreted factor genes’ mRNA levels in sorted macrophages, normalized to *rp49*, at 96 h AEL. n=3-4, each sample pooled of 20,000 macrophages. (D) Depletion of *spi, dpp, gbb, upd2, upd3, dilp6, pvf1, pvf3, akh, idgf2, idgf6*, and *spz* have no impact on the developmental timing of the larva to pupa transition. Median times of puparium formation (hrs AEL) were analyzed by a two-tailed unpaired *t-*test. n.s (no statistical significances).

**Figure S3.**
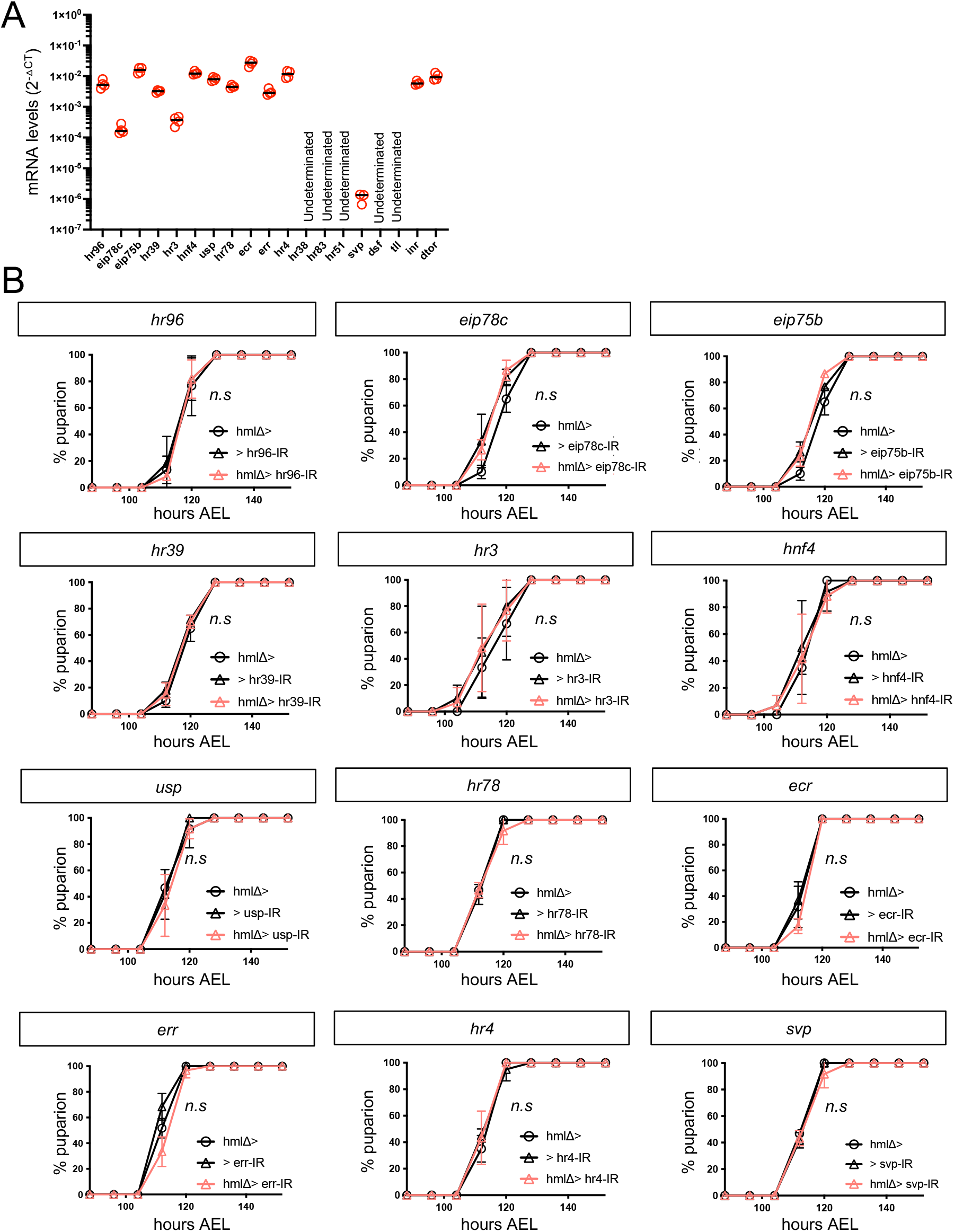
Screen for nutrient receptors expressed by larval hemocytes. (A) Candidate nutritional sensor genes mRNA levels in macrophages normalized to *rp49* at 96 h AEL of sorted macrophages. n=3-4, each sample pooled of 20000 macrophages. (B) Depletion of *hr96, eip78c, eip75b, hr39, hr3, hnf4, usp, hr78, ecr, err, hr4*, and *svp* have not any impact in the developmental timing program in the larva top pupa transition. Median time of puparium formation (hours AEL) were analyzed by a two-tailed unpaired *t-*test. n.s (no statistical significances).

**Figure S4.**
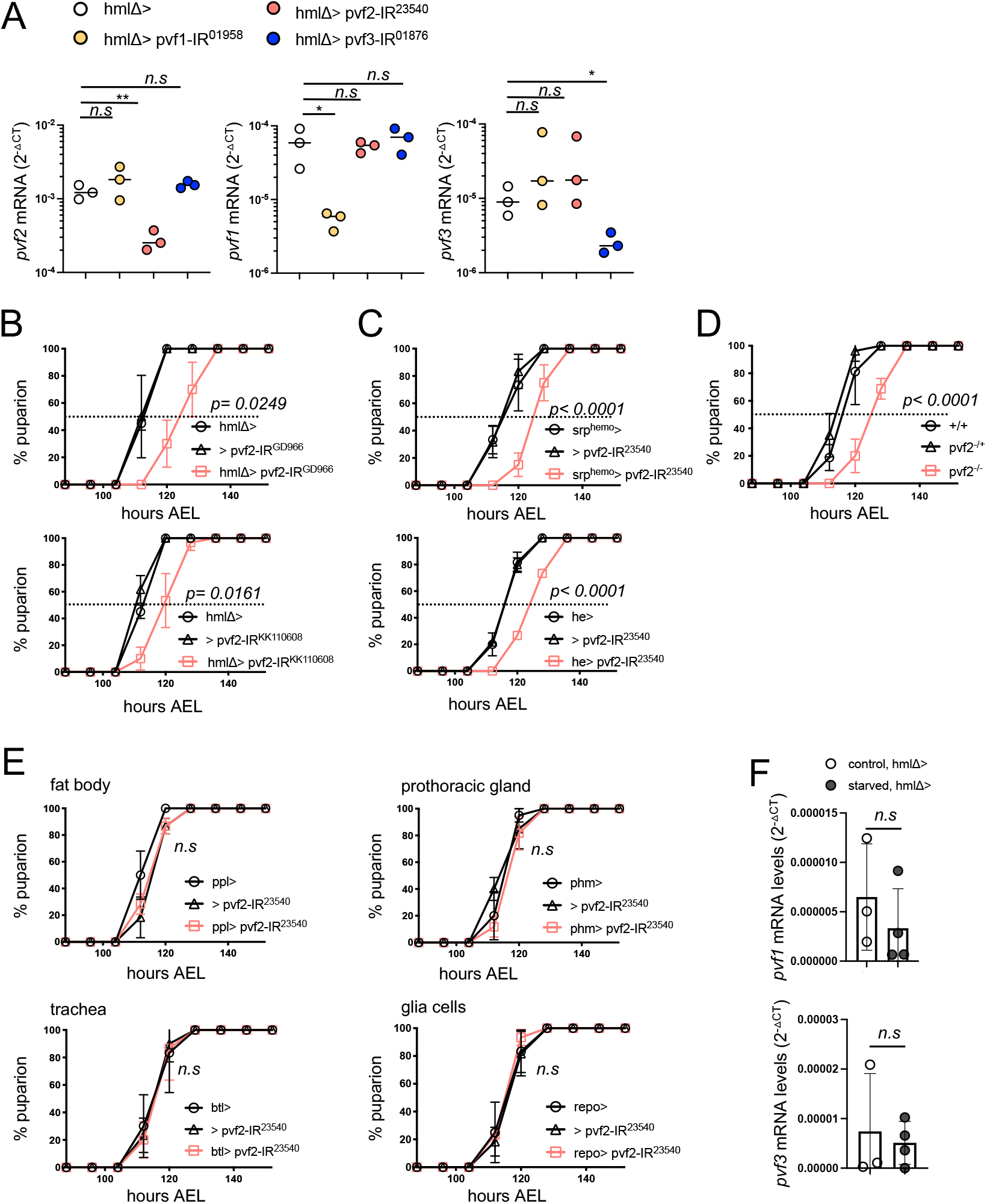
Production of Pvf2 by macrophages is required and sufficient for developmental timing by regulating ecdysone production. (A) RNAi efficiency of *pvf1, pvf2*, and *pvf3* RNAi lines from BDSC. mRNA levels normalized to *rp49* at 96 h AEL of sorted macrophages in the genotypes indicated. n=3, each sample pooled of 20,000 macrophages. (B) Knock-down of *pvf2*^*GD966*^ and *pvf2*^KK110608 specifically^ in macrophages does not induce developmental delay (n=3, 20 larvae per genotype in 3 independent crosses, 60 larvae in total per condition). (C) Knock-down of *pvf2* specifically in macrophages, using *srp*^*hemo*^*-gal4* and *he-gal4*, induces developmental delay (n=3, 20 larvae per genotype in 3 independent crosses, 60 larvae in total per condition). (D) *pvf2*^*c06947*^ loss-of-function flies show a developmental delay in the larva-to-pupa transition (n=4, 20 larvae per genotype in 4 independent crosses, 80 larvae in total per condition). (E) Knock-down of *pvf2* specifically in fat body (*ppl-gal4* driver), prothoracic gland (*phm-gal4* driver), trachea (*btl-gal4* driver), and glia cells (*repo-gal4* driver) (n=3, 20 larvae per genotype in 3 independent crosses, 60 larvae in total per condition). (F) mRNA levels of the pvf secreted factors from macrophages, *pvf1*,and *pvf3* in sorted GFP^+^ macrophages in the starved larvae at 96 hours AEL. mRNA levels are normalized to *rp49*. n=3-4, each sample a pool of 20,000 GFP^+^ macrophages. Data were analyzed by a two-tailed unpaired *t-*test and values represent the mean ± SD. *p < 0.05, **p < 0.01, n.s (not significant).

**Figure S5.**
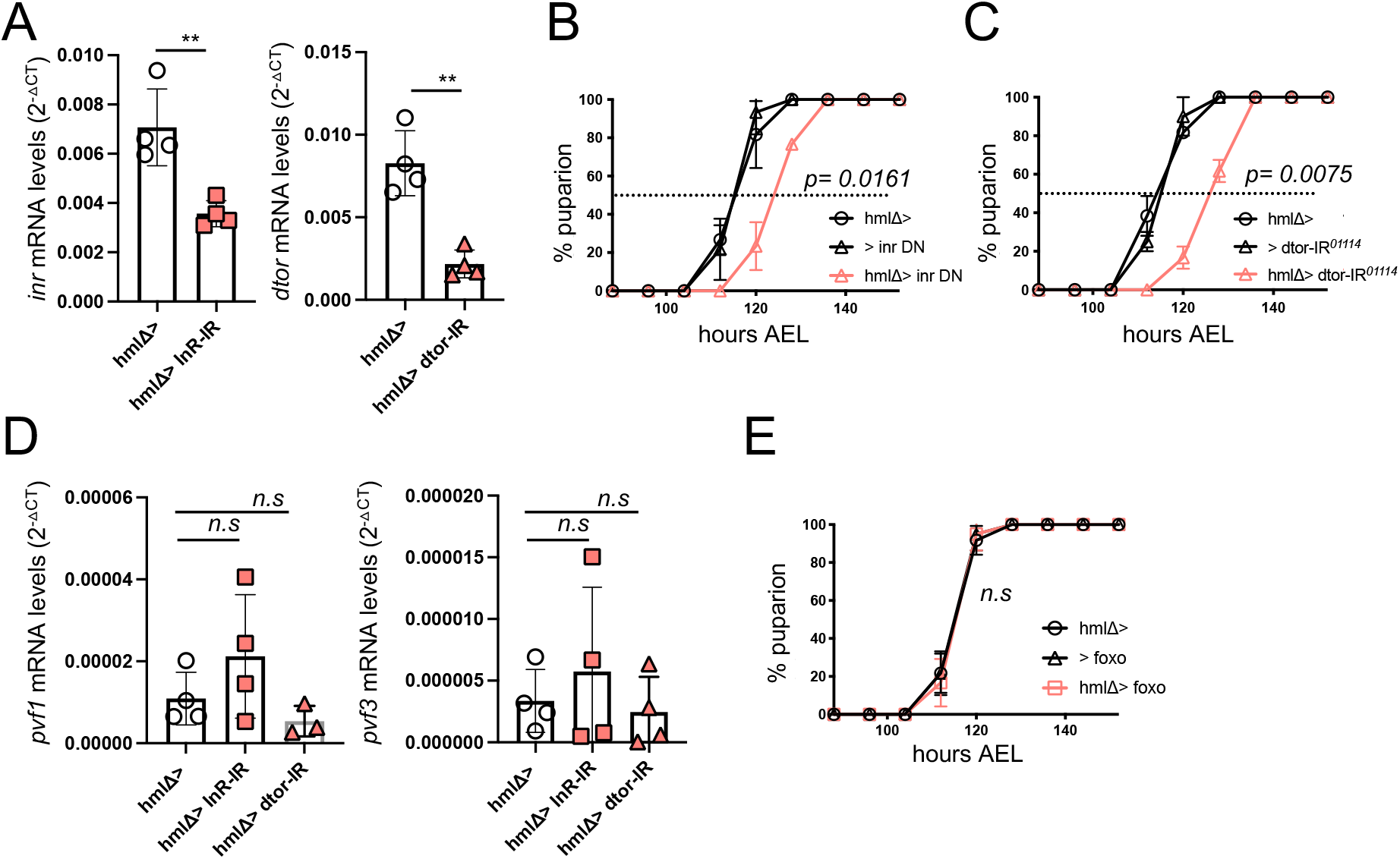
Inr and dTor control production of Pvf2 by macrophages. (A) RNAi efficiency of BDSC lines for *inr* and *dtor* in sorted GFP^+^ macrophages from the indicated genotype at 96 hours AEL. mRNA levels are normalized to *rp49*. n=4, 20,000 GFP^+^ macrophages pooled for each sample. (B) Overexpression specifically in macrophages of the dominant negative isoform of *inr* induces developmental delay (n=3, 20 larvae per genotype in 3 independent crosses, 60 larvae in total per condition). (C) Knock-down of *dtor* specifically in macrophages induces developmental delay (n=3, 20 larvae per genotype in 3 independent crosses, 60 larvae in total per condition). (D) mRNA levels of the secreted Pvf factors *pvf1* and *pvf3* in sorted GFP^+^ macrophages from the indicated genotype at 96 hours AEL. mRNA levels are normalized to *rp49*. n=4, each sample pooled of 20000 GFP^+^ macrophages. (E) Overexpression of *foxo* specifically in macrophages does not induces developmental delay (n=3, 20 larvae per genotype in 3 independent crosses, 60 larvae in total per condition). Data were analyzed by a two-tailed unpaired *t-*test and values represent the mean ± SD., **p < 0.01,. n.s (not significant).

**Figure S6.**
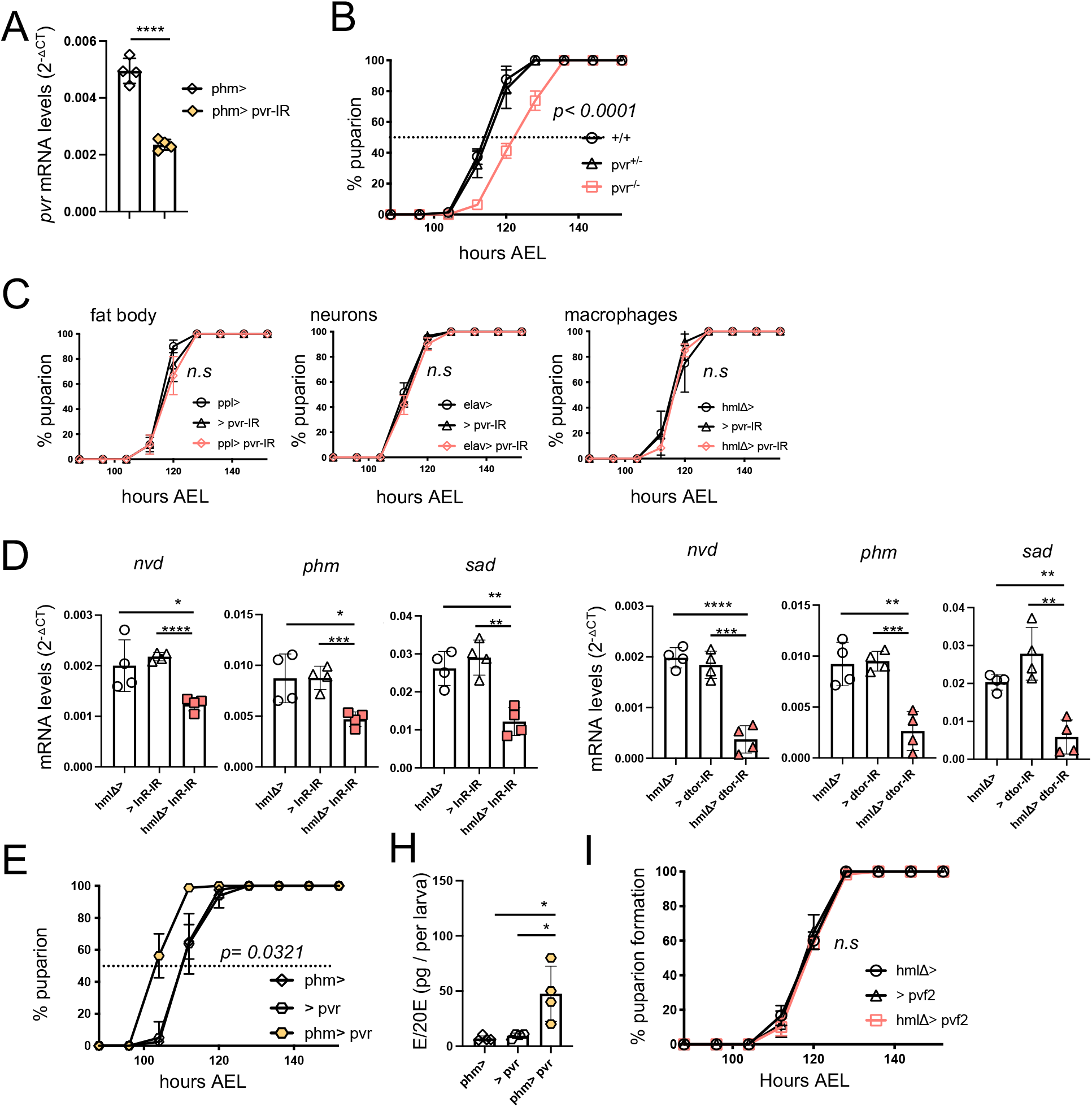
Macrophage’s *inr/dtor/pvf2* genetic cassette mediates ecdysone synthesis by *pvr* in the prothoracic gland. (A) mRNA levels of *pvr* in brain-ring gland complex at 96 hours AEL. mRNA levels are normalized to *rp49*. n=4, each sample pooled of 5-7 brain-ring gland complex. (B) *pvr*^*1*^ hypomorphic mutant flies show developmental delay in the larva-to-pupa transition (n=4, 20 larvae per genotype in 4 independent crosses, 80 larvae in total per condition). (C) knock-down of *pvr* specifically in fat body (*ppl-gal4* driver), neurons (*elav-gal4* driver), and macrophages (*hmlΔ-gal4*) (n=3, 20 larvae per genotype in 3 independent crosses, 60 larvae in total per condition). (D) Transcriptional levels of the ecdysone biosynthesis gene *nvd, phm, sad, dib*, and *spo* in *inr* and *dtor* knock down in macrophage larvae. mRNA levels normalized to *rp49* at 104 hours AEL. n=4. Sample were pooled by 5 larvae per condition and genotype. (E) Overexpression of *pvr* specifically in the prothoracic gland induces acceleration of developmental timing (n=3, 20 larvae per genotype in 3 independent crosses, 60 larvae in total per condition). (H) Ecdysone total levels in whole larvae in the indicate genotype at 96 hours AEL, quantified by ELISA. n=4, each sample pooled of 10 larvae. (I) *pvf2* overexpression specifically in macrophages, using *hemolectin*Δ*-gal4*, does not induce developmental acceleration (n=3, 20 larvae per genotype in 3 independent crosses, 60 larvae in total per condition). Data were analyzed by a two-tailed unpaired *t-*test and values represent the mean ± SD. *p < 0.05, **p < 0.01, ***p < 0.001, ****p < 0.0001. n.s (not significant).

**Figure S7.**
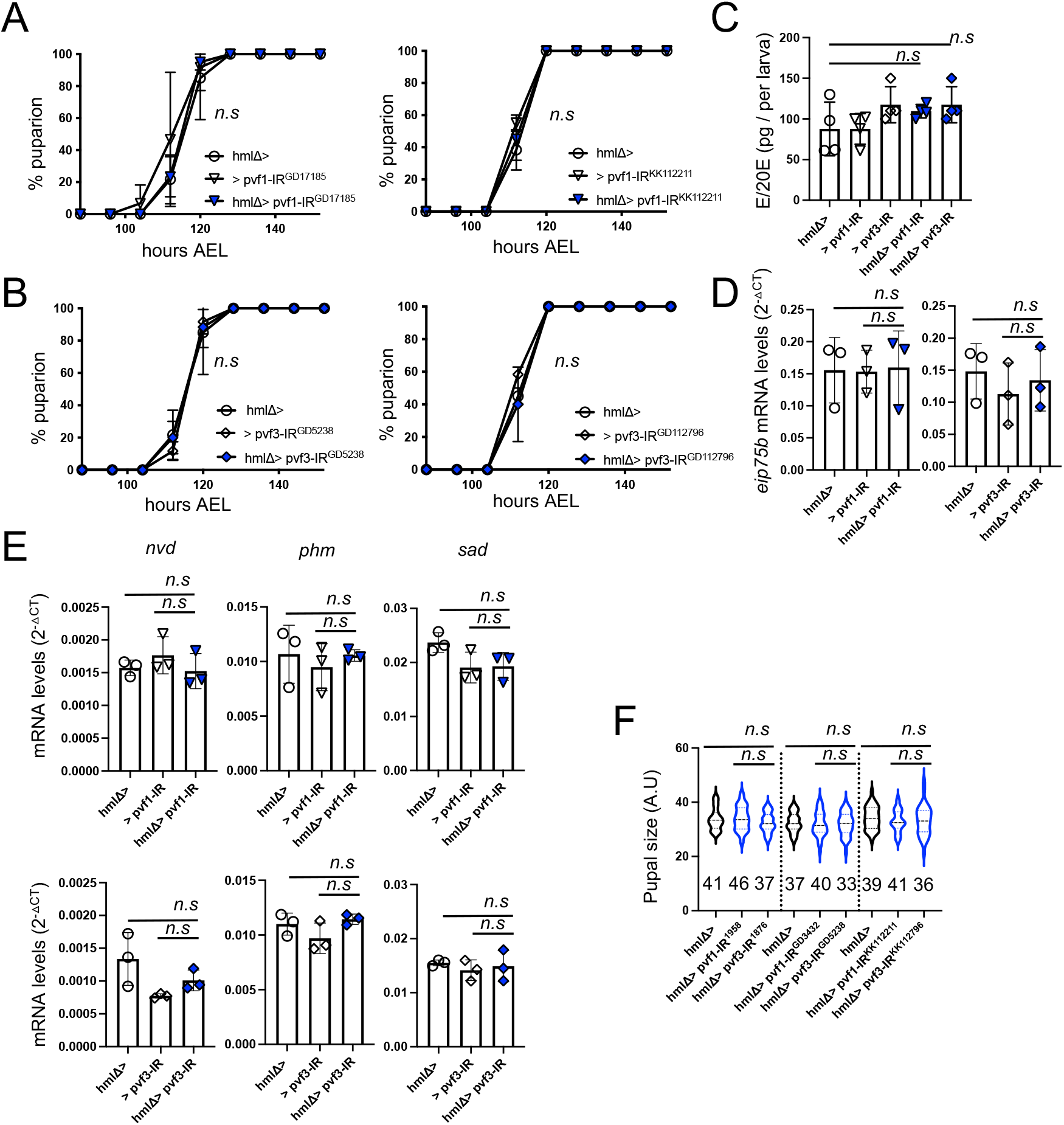
Macrophage-derived Pvf1 and Pvf3 does not regulate ecdysone synthesis and developmental timing. (A) knock-down of *pvf1* ^GD17185^ and *pvf1* ^KK112211^ RNAi lines specifically in macrophages does not induce developmental delay (n=3, 20 larvae per genotype in 3 independent crosses, 60 larvae in total per condition). (B) knock-down of *pvf3* ^GD5238^ and *pvf3* ^KK112796^ specifically in macrophages does not induce developmental delay (n=3, 20 larvae per genotype in 3 independent crosses, 60 larvae in total per condition). (C) Whole larvae ecdysone total levels in *pvf1* and *pvf3* RNAi from BDSC knock down in macrophages, quantified by ELISA. n=4, each sample pooled of 10 larvae. (D) Transcriptional levels of target gene for ecdysone signaling *eip75b* in *pvf1* and *pvf3* knock down in macrophages. mRNA levels normalized to *rp49* at 104 hours AEL. (E) Transcriptional levels of target gene for ecdysone signaling *eip75b* in *pvf1* and *pvf3* knock down in macrophages. mRNA levels normalized to *rp49* at 104 hours AEL. n=3. Each sample pooled of 5 whole larvae at each time point and genotype indicated. (F) Final pupal size in *pvf1* and *pvf3* knock down in macrophages. Data were analyzed by a two-tailed unpaired *t-*test and values represent the mean ± SD. n.s (not significant).

## References

1. Poethig, R.S., Phase change and the regulation of developmental timing in plants. Science, 2003. 301(5631): p. 334–6.

2. Plant, T.M., Neuroendocrine control of the onset of puberty. Front Neuroendocrinol, 2015. 38: p. 73–88.

3. Bordini, B. and R.L. Rosenfield, Normal pubertal development: Part I: The endocrine basis of puberty. Pediatr Rev, 2011. 32(6): p. 223–9.

4. McBrayer, Z., et al., Prothoracicotropic hormone regulates developmental timing and body size in Drosophila. Dev Cell, 2007. 13(6): p. 857–71.

5. Rewitz, K.F., et al., The insect neuropeptide PTTH activates receptor tyrosine kinase torso to initiate metamorphosis. Science, 2009. 326(5958): p. 1403–5.

6. Herbison, A.E., Control of puberty onset and fertility by gonadotropin-releasing hormone neurons. Nat Rev Endocrinol, 2016. 12(8): p. 452–66.

7. Yamanaka, N., K.F. Rewitz, and M.B. O’Connor, Ecdysone control of developmental transitions: lessons from Drosophila research. Annu Rev Entomol, 2013. 58: p. 497–516.

8. Rewitz, K.F., N. Yamanaka, and M.B. O’Connor, Developmental checkpoints and feedback circuits time insect maturation. Curr Top Dev Biol, 2013. 103: p. 1–33.

9. Imura, E., et al., The Corazonin-PTTH Neuronal Axis Controls Systemic Body Growth by Regulating Basal Ecdysteroid Biosynthesis in Drosophila melanogaster. Curr Biol, 2020. 30(11): p. 2156–2165 e5.

10. Bateman, J.M. and H. McNeill, Temporal control of differentiation by the insulin receptor/tor pathway in Drosophila. Cell, 2004. 119(1): p. 87–96.

11. Villamor, E. and E.C. Jansen, Nutritional Determinants of the Timing of Puberty. Annu Rev Public Health, 2016. 37: p. 33–46.

12. Danielsen, E.T., M.E. Moeller, and K.F. Rewitz, Nutrient signaling and developmental timing of maturation. Curr Top Dev Biol, 2013. 105: p. 37–67.

13. Layalle, S., N. Arquier, and P. Leopold, The TOR pathway couples nutrition and developmental timing in Drosophila. Dev Cell, 2008. 15(4): p. 568–77.

14. Tanner, J.M., Trend towards earlier menarche in London, Olso, Copenhagen, the Netherlands and Hungary. Nature, 1973. 243(5402): p. 95–6.

15. Zacharias, L. and R.J. Wurtman, Age at menarche. Genetic and environmentalinfluences. N Engl J Med, 1969. 280(16): p. 868–75.

16. Marshall, J.C. and R.P. Kelch, Low dose pulsatile gonadotropin-releasing hormone in anorexia nervosa: a model of human pubertal development. J Clin Endocrinol Metab, 1979. 49(5): p. 712–8.

17. Juarez-Carreno, S., et al., Body-fat sensor triggers ribosome maturation in the steroidogenic gland to initiate sexual maturation in Drosophila. Cell Rep, 2021. 37(2): p. 109830.

18. Okabe, Y. and R. Medzhitov, Tissue biology perspective on macrophages. Nat Immunol, 2016. 17(1): p. 9–17.

19. Gold, K.S. and K. Bruckner, Macrophages and cellular immunity in Drosophila melanogaster. Semin Immunol, 2015. 27(6): p. 357–68.

20. Wood, W. and P. Martin, Macrophage Functions in Tissue Patterning and Disease: New Insights from the Fly. Dev Cell, 2017. 40(3): p. 221–233.

21. Rankin, L.C. and D. Artis, Beyond Host Defense: Emerging Functions of the Immune System in Regulating Complex Tissue Physiology. Cell, 2018. 173(3): p. 554–567.

22. Cox, N., et al., Origins, Biology, and Diseases of Tissue Macrophages. Annu Rev Immunol, 2021. 39: p. 313–344.

23. Cox, N., et al., Diet-regulated production of PDGFcc by macrophages controls energy storage. Science, 2021. 373(6550).

24. Woodcock, K.J., et al., Macrophage-derived upd3 cytokine causes impaired glucose homeostasis and reduced lifespan in Drosophila fed a lipid-rich diet. Immunity, 2015. 42(1): p. 133–44.

25. Chen, T.T., et al., The effect of peritoneal macrophage-derived factor(s) on ovarian progesterone secretion and LH receptors: the role of calcium. Am J Reprod Immunol, 1992. 28(1): p. 43–50.

26. Gu, X.W., et al., Immune Cells as Critical Regulators of Steroidogenesis in the Testis and Beyond. Frontiers in Endocrinology, 2022. 13.

27. Shin, M., et al., Subpopulation of Macrophage-Like Plasmatocytes Attenuates Systemic Growth via JAK/STAT in the Drosophila Fat Body. Front Immunol, 2020. 11: p. 63.

28. Jacome-Galarza, C.E., et al., Developmental origin, functional maintenance and genetic rescue of osteoclasts. Nature, 2019. 568(7753): p. 541–545.

29. Cohen, P.E., et al., Macrophages: important accessory cells for reproductive function. J Leukoc Biol, 1999. 66(5): p. 765–72.

30. Lokka, E., et al., Generation, localization and functions of macrophages during the development of testis. Nat Commun, 2020. 11(1): p. 4375.

31. Norman, R.J. and M. Brannstrom, White cells and the ovary--incidental invaders or essential effectors? J Endocrinol, 1994. 140(3): p. 333–6.

32. Zhang, Z., L. Huang, and L. Brayboy, Macrophages: an indispensable piece of ovarian health. Biol Reprod, 2021. 104(3): p. 527–538.

33. Cohen, P.E., et al., Colony-stimulating factor 1 regulation of neuroendocrine pathways that control gonadal function in mice. Endocrinology, 2002. 143(4): p. 1413–22.

34. Cohen, P.E., M.P. Hardy, and J.W. Pollard, Colony-stimulating factor-1 plays a major role in the development of reproductive function in male mice. Mol Endocrinol, 1997. 11(11): p. 1636–50.

35. Cohen, P.E., et al., Absence of colony-stimulating factor-1 in osteopetrotic (csfmop/csfmop) mice results in male fertility defects. Biol Reprod, 1996. 55(2): p. 310–7.

36. Yee, J.B. and J.C. Hutson, Effects of testicular macrophage-conditioned medium on Leydig cells in culture. Endocrinology, 1985. 116(6): p. 2682–4.

37. Matzuk, M.M. and D.J. Lamb, The biology of infertility: research advances and clinical challenges. Nat Med, 2008. 14(11): p. 1197–213.

38. Ohhara, Y., S. Kobayashi, and N. Yamanaka, Nutrient-Dependent Endocycling in Steroidogenic Tissue Dictates Timing of Metamorphosis in Drosophila melanogaster. PLoS Genet, 2017. 13(1): p. e1006583.

39. Koyama, T., et al., Nutritional control of body size through FoxO-Ultraspiracle mediated ecdysone biosynthesis. Elife, 2014. 3.

40. Segraves, W.A. and D.S. Hogness, The E75 ecdysone-inducible gene responsible for the 75B early puff in Drosophila encodes two new members of the steroid receptor superfamily. Genes Dev, 1990. 4(2): p. 204–19.

41. Charroux, B. and J. Royet, Elimination of plasmatocytes by targeted apoptosis reveals their role in multiple aspects of the Drosophila immune response. Proc Natl Acad Sci U S A, 2009. 106(24): p. 9797–802.

42. Stephenson, H.N., et al., Hemocytes are essential for Drosophila melanogaster post-embryonic development, independent of control of the microbiota. Development, 2022. 149(18).

43. Gilbert, L.I., R. Rybczynski, and J.T. Warren, Control and biochemical nature of the ecdysteroidogenic pathway. Annu Rev Entomol, 2002. 47: p. 883–916.

44. Mase, A., J. Augsburger, and K. Bruckner, Macrophages and Their Organ Locations Shape Each Other in Development and Homeostasis - A Drosophila Perspective. Front Cell Dev Biol, 2021. 9: p. 630272.

45. Belyaeva, V., et al., Fos regulates macrophage infiltration against surrounding tissue resistance by a cortical actin-based mechanism in Drosophila. PLoS Biol, 2022. 20(1): p. e3001494.

46. Cho, N.K., et al., Developmental control of blood cell migration by the Drosophila VEGF pathway. Cell, 2002. 108(6): p. 865–76.

47. Boucher, J., A. Kleinridders, and C.R. Kahn, Insulin receptor signaling in normal and insulin-resistant states. Cold Spring Harb Perspect Biol, 2014. 6(1).

48. Saxton, R.A. and D.M. Sabatini, mTOR Signaling in Growth, Metabolism, and Disease. Cell, 2017. 169(2): p. 361–371.

49. Saltiel, A.R. and C.R. Kahn, Insulin signalling and the regulation of glucose and lipid metabolism. Nature, 2001. 414(6865): p. 799–806.

50. Britton, J.S., et al., Drosophila’s insulin/PI3-kinase pathway coordinates cellular metabolism with nutritional conditions. Dev Cell, 2002. 2(2): p. 239–49.

51. Shingleton, A.W., et al., The temporal requirements for insulin signaling during development in Drosophila. PLoS Biol, 2005. 3(9): p. e289.

52. Grewal, S.S., Insulin/TOR signaling in growth and homeostasis: a view from the fly world. Int J Biochem Cell Biol, 2009. 41(5): p. 1006–10.

53. Pan, X. and M.B. O’Connor, Coordination among multiple receptor tyrosine kinase signals controls Drosophila developmental timing and body size. Cell Rep, 2021. 36(9): p. 109644.

54. Bruckner, K., et al., The PDGF/VEGF receptor controls blood cell survival in Drosophila. Dev Cell, 2004. 7(1): p. 73–84.

55. Zheng, H., et al., Premature remodeling of fat body and fat mobilization triggered by platelet-derived growth factor/VEGF receptor in Drosophila. FASEB J, 2017. 31(5): p. 1964–1975.

56. Read, R.D., Pvr receptor tyrosine kinase signaling promotes post-embryonic morphogenesis, and survival of glia and neural progenitor cells in Drosophila. Development, 2018. 145(23).

57. Munier, A.I., et al., PVF2, a PDGF/VEGF-like growth factor, induces hemocyte proliferation in Drosophila larvae. EMBO Rep, 2002. 3(12): p. 1195–200.

58. Tattikota, S.G., et al., A single-cell survey of Drosophila blood. Elife, 2020. 9.

59. Colombani, J., et al., Antagonistic actions of ecdysone and insulins determine final size in Drosophila. Science, 2005. 310(5748): p. 667–70.

60. Mirth, C., J.W. Truman, and L.M. Riddiford, The role of the prothoracic gland in determining critical weight for metamorphosis in Drosophila melanogaster. Curr Biol, 2005. 15(20): p. 1796–807.

61. Okamoto, N. and N. Yamanaka, Nutrition-dependent control of insect development by insulin-like peptides. Curr Opin Insect Sci, 2015. 11: p. 21–30.

62. Delanoue, R., et al., Drosophila insulin release is triggered by adipose Stunted ligand to brain Methuselah receptor. Science, 2016. 353(6307): p. 1553–1556.

63. Geminard, C., E.J. Rulifson, and P. Leopold, Remote control of insulin secretion by fat cells in Drosophila. Cell Metab, 2009. 10(3): p. 199–207.

64. Rajan, A. and N. Perrimon, Drosophila cytokine unpaired 2 regulates physiological homeostasis by remotely controlling insulin secretion. Cell, 2012. 151(1): p. 123–37.

65. Agrawal, N., et al., The Drosophila TNF Eiger Is an Adipokine that Acts on Insulin-Producing Cells to Mediate Nutrient Response. Cell Metab, 2016. 23(4): p. 675–84.

66. Ratheesh, A., V. Belyaeva, and D.E. Siekhaus, Drosophila immune cell migration and adhesion during embryonic development and larval immune responses. Curr Opin Cell Biol, 2015. 36: p. 71–9.

67. Parsons, B. and E. Foley, The Drosophila platelet-derived growth factor and vascular endothelial growth factor-receptor related (Pvr) protein ligands Pvf2 and Pvf3 control hemocyte viability and invasive migration. J Biol Chem, 2013. 288(28): p. 20173–83.

68. Droujinine, I.A. and N. Perrimon, Interorgan Communication Pathways in Physiology: Focus on Drosophila. Annu Rev Genet, 2016. 50: p. 539–570.

69. Gnessi, L., et al., Rat Leydig cells bind platelet-derived growth factor through specific receptors and produce platelet-derived growth factor-like molecules. Endocrinology, 1992. 130(4): p. 2219–24.

70. Gnessi, L., et al., Testicular development involves the spatiotemporal control of PDGFs and PDGF receptors gene expression and action. J Cell Biol, 1995. 131(4): p. 1105–21.

71. Loveland, K.L., et al., Platelet-derived growth factor ligand and receptor subunit mRNA in the Sertoli and Leydig cells of the rat testis. Mol Cell Endocrinol, 1995. 108(1-2): p. 155–9.

72. Taylor, C.C., Platelet-derived growth factor activates porcine thecal cell phosphatidylinositol-3-kinase-Akt/PKB and ras-extracellular signal-regulated kinase-1/2 kinase signaling pathways via the platelet-derived growth factor-beta receptor. Endocrinology, 2000. 141(4): p. 1545–53.

73. Basciani, S., et al., Expression of platelet-derived growth factor-A (PDGF-A), PDGF-B, and PDGF receptor-alpha and -beta during human testicular development and disease. J Clin Endocrinol Metab, 2002. 87(5): p. 2310–9.

74. Brennan, J., C. Tilmann, and B. Capel, Pdgfr-alpha mediates testis cord organization and fetal Leydig cell development in the XY gonad. Genes Dev, 2003. 17(6): p. 800–10.

75. Soriano, P., Abnormal kidney development and hematological disorders in PDGF beta-receptor mutant mice. Genes Dev, 1994. 8(16): p. 1888–96.

76. Soriano, P., The PDGF alpha receptor is required for neural crest cell development and for normal patterning of the somites. Development, 1997. 124(14): p. 2691–700.

77. Schmahl, J., K. Rizzolo, and P. Soriano, The PDGF signaling pathway controls multiple steroid-producing lineages. Genes Dev, 2008. 22(23): p. 3255–67.

## Supplementary references

1. Sinenko, S.A. and B. Mathey-Prevot, Increased expression of Drosophila tetraspanin, Tsp68C, suppresses the abnormal proliferation of ytr-deficient and Ras/Raf-activated hemocytes. Oncogene, 2004. 23(56): p. 9120–8.

2. Ramond, E., et al., The adipokine NimrodB5 regulates peripheral hematopoiesis in Drosophila. FEBS J, 2020. 287(16): p. 3399–3426.

3. Colombani, J., et al., A nutrient sensor mechanism controls Drosophila growth. Cell, 2003. 114(6): p. 739–49.

4. Rulifson, E.J., S.K. Kim, and R. Nusse, Ablation of insulin-producing neurons in flies: growth and diabetic phenotypes. Science, 2002. 296(5570): p. 1118–20.

5. Palomino-Schatzlein, M., et al., A toolbox to study metabolic status of Drosophila melanogaster larvae. STAR Protoc, 2022. 3(1): p. 101195.

6. Rosin, D., et al., Apical accumulation of the Drosophila PDGF/VEGF receptor ligands provides a mechanism for triggering localized actin polymerization. Development, 2004. 131(9): p. 1939–48.

7. Texada, M.J., et al., Autophagy-Mediated Cholesterol Trafficking Controls Steroid Production. Dev Cell, 2019. 48(5): p. 659–671 e4.

8. Ikeya, T., et al., Nutrient-dependent expression of insulin-like peptides from neuroendocrine cells in the CNS contributes to growth regulation in Drosophila. Curr Biol, 2002. 12(15): p. 1293–300.

9. Tattikota, S.G., et al., A single-cell survey of Drosophila blood. Elife, 2020. 9.

